# A Sense of Place: Transcriptomics Identifies Environmental Signatures in Cabernet Sauvignon Berry Skins in the Late Stages of Ripening

**DOI:** 10.1101/729236

**Authors:** Grant R. Cramer, Noé Cochetel, Ryan Ghan, Agnès Destrac-Irvine, Serge Delrot

**Affiliations:** Department of Biochemistry and Molecular Biology, University of Nevada, Reno, Reno, NV 89557, USA; UMR Ecophysiology and Grape Functional Genomics, Institut des Sciences de la Vigne et du Vin, University of Bordeaux, Villenave d’Ornon, France

**Keywords:** abiotic stress, biotic stress, grape berry development, RNA-Seq, transcriptomics, *Vitis vinifera* L

## Abstract

**Background:** Grape berry ripening is influenced by climate, the main component of the “terroir” of a place. Light and temperature are major factors in the vineyard that affect berry development and fruit metabolite composition.

**Results:** To better understand the effect of “place” on transcript abundance during the late stages of berry ripening, Cabernet Sauvignon berries grown in Bordeaux and Reno were compared at similar sugar levels (19 to 26 °Brix (total soluble solids)). Day temperatures were warmer and night temperatures were cooler in Reno. °Brix was lower in Bordeaux berries compared to Reno at maturity levels considered optimum for harvest. RNA-Seq analysis identified 5528 differentially expressed genes between Bordeaux and Reno grape skins at 22°Brix. Weighted Gene Coexpression Network Analysis for all expressed transcripts for all four °Brix levels measured indicated that the majority (75%) of transcript expression differed significantly between the two locations. Top gene ontology categories for the common transcript sets were translation, photosynthesis, DNA metabolism and catabolism. Top gene ontology categories for the differentially expressed genes at 22°Brix involved response to stimulus, biosynthesis and response to stress. Some differentially expressed genes encoded terpene synthases, cell wall enzymes, kinases, transporters, transcription factors and photoreceptors. Most circadian clock genes had higher transcript abundance in Bordeaux. Bordeaux berries had higher transcript abundance with differentially expressed genes associated with seed dormancy, light, auxin, ethylene signaling, powdery mildew infection, phenylpropanoid, carotenoid and terpenoid metabolism, whereas Reno berries were enriched with differentially expressed genes involved in water deprivation, cold response, ABA signaling and iron homeostasis.

**Conclusions:** Transcript abundance profiles in the berry skins at maturity were highly dynamic. RNA-Seq analysis identified a smaller (25% of total) common core set of ripening genes that appear not to depend on rootstock, vineyard management, plant age, soil and climatic conditions. Much of the gene expression differed between the two locations and could be associated with multiple differences in environmental conditions that may have affected the berries in the two locations; some of these genes may be potentially controlled in different ways by the vinegrower to adjust final berry composition and reach a desired result.

## Background

*Vitis vinifera* grapevines originated approximately 65 million years ago from Eurasia and have been cultivated for at least the last 8000 years for its fruits that are crushed to make wine [1]. Grapevines are now grown throughout the world in many kinds of environments.

Grape berry development is a complex process involving three developmental phases and multiple hormones [2, 3]. It is in the latter ripening phase that many compounds involved in flavor and aromas are synthesized, conjugated or catabolized. Most of these compounds reside in the skin of the berry and seem to develop in the very last stages of berry development [4–6]. Aroma and flavor are important sensory components of wine. They are derived from multiple classes of compounds in grapes including important volatile compounds from the grape and from yeast metabolism during grape fermentation [5, 6]. Each grape cultivar produces a unique set of volatile and flavor compounds at varying concentration that represents its wine typicity or typical cultivar characteristics [6]. Esters and terpenes are volatile compound chemical classes largely responsible for the fruity and floral aromas in wines [5, 6]. Esters are largely produced during yeast fermentation from grape-derived products such as aliphatic alcohols and aldehydes [7, 8]. Grape lipoxygenases are thought to provide the six carbon precursors from fatty acids for the synthesis of the fruity aroma, hexyl acetate [8], in yeast during wine fermentation. Terpenes mostly originate from the grapes and are found in both the free and bound (glycosylated) forms. Both plant fatty acid and terpenoid metabolism pathways are very sensitive to the environment [9–13]. Climate has large effects on berry development and composition [14–16]. Besides grape genetics other factors may influence metabolite composition including the local grape berry microbiome [17], the soil type [15] and the rootstock [18–22]. While there is evidence that rootstock can affect fruit composition and transcript abundance, this effect appears to be minor relative to other environmental factors [18, 19, 21]. Many cultural practices used by the grape grower may directly or indirectly affect the environment sensed by the grapevine (row orientation, planting density, pruning, leaf removal, etc.). Temperature and light are major contributors to “terroir”. The terroir term is used in the wine industry to acknowledge the influence of the environment on the grapevine and the distinctive characteristics that are contributed to the typicity of a wine [2, 14, 15, 23]. It includes the biotic, abiotic and soil environments as well as the viticultural practices. In the present work, we will use the term “place” to address all of the above except for the viticultural practices.

Recently, a transcriptomic approach was used to elucidate the common gene subnetworks of the late stages of berry development when grapes are normally harvested at their peak maturity [4]. One of the major subnetworks associated with ripening involved autophagy, catabolism, RNA splicing, proteolysis, chromosome organization and the circadian clock. An integrated model was constructed to link light sensing with the circadian clock highlighting the importance of the light environment on berry development. In this report, in order to get a better understanding of how much of the gene expression in Cabernet Sauvignon berry skin could be attributed to environmental influences, we tested the hypothesis that there would be significant differences in gene expression during the late stages of Cabernet Sauvignon berry ripening between two widely different locations: one in Reno, NV, USA (RNO) and the other in Bordeaux, France (BOD). The analysis revealed a core set of genes that did not depend on location, climate, vineyard management, grafting and soil properties. Also, the analysis revealed key genes that are differentially expressed between the two locations and links some of these differences to the effects of temperature and other environmental factors on aromatic and other quality-trait-associated pathways. Many gene families were differentially expressed and may provide useful levers for the vinegrower to adjust berry composition. Among others, these families encompassed genes involved in amino acid and phenylpropanoid metabolism, as well as aroma and flavor synthesis.

## Results

### Background Data for a “Sense of Place”

To test the hypothesis that the transcript abundance of grape berries during the late stages of ripening differed in two locations with widely different environmental conditions, we compared the transcript abundance of grape berry skins in BOD and RNO. The vineyards were planted in 2004 and 2009 in RNO and BOD, respectively. BOD vines were grafted on to SO4 rootstock and RNO vines were grown on their own roots. Both BOD and RNO used a vertical shoot positioning trellis design. The environmental variables between the two vineyard sites had a number of differences. BOD has a slightly more northern latitude than RNO; consequently, day lengths were slightly longer in BOD at the beginning of harvest and slightly shorter at the end of harvest (Table 1). On the final harvest dates, the day length differed between the two locations by about 30 min.

**Table 1.** Environmental variables for the harvest times in BOD and RNO. Grapes were harvested in RNO from September 10 to 26, 2012 and in BOD from September 17 to October 8, 2013.

| Environmental Variable | BOD (2013) | RNO (2012) |
| --- | --- | --- |
| Elevation (m) | 25 | 1,373 |
| Average Daily Solar Radiation (kW-hr m<sup>-2</sup>) | 1.17 | 5.86 |
| Day length Starting Harvest Date | 12:25:36 | 12:38:34 |
| Day length Ending Harvest Date | 11:20:57 | 11:57:37 |
| Maximum Temperature Starting Harvest Date | 19 | 30.5 |
| Minimum Temperature Starting Harvest Date | 13 | 13.9 |
| Maximum Temperature Ending Harvest Date | 18 | 27.8 |
| Minimum Temperature Ending Harvest Date (°C) | 11 | 6.7 |
| Ave September Maximum Temperature (°C) | 23.9 | 30.2 |
| Ave September Minimum Temperature (°C) | 13.9 | 10.2 |
| Latitude | 44°47'23.83" N | 39°52'96" N |
| Longitude | 0°34'39.3" W | 119°81'38" W |
| September Precipitation (mm) | 65.5 | 2.03 |
| Average Monthly Relative Humidity (%) | 74 | 34 |
| Soil Type | Gravelly soil | Sandy Loam |
| Soil pH | 6.2 | 6.7 |
| Soil Fe (µg g<sup>-1</sup>) | 193 | 38 |
| Root stock | SO4 | Own-rooted |

Average monthly maximum temperatures were warmer in RNO than BOD, but minimum September temperatures were cooler (Table 1). This made for an average daily day/night temperature differential of 10°C and 20°C during the harvest periods for BOD and RNO, respectively. The day temperatures were warmer by about 6°C and the night time temperatures were about 4°C cooler in RNO.

The RNO site was much drier than the BOD site (Table 1). September monthly precipitation totals were 65.5 and 2.03 mm and had average relative humidities of 74 and 34% for BOD and RNO, respectively. The soil at the BOD site was a gravelly soil with a pH of 6.2 and the soil at the RNO site was a deep sandy loam with a pH of 6.7. No pathogen, nutrient deficiency or toxicity symptoms were evident in the vines.

### Transcriptomics

The analysis of transcript profiles of Cabernet Sauvignon grapes harvested in RNO in September of 2012 was previously described [4]. Individual berry skins were separated immediately from the whole berry and the individual total soluble solids (°Brix) level of the berry, which is mostly composed of sugars, was determined. Similarly, Cabernet Sauvignon berry skins from BOD were harvested from the middle of September in 2013 until the end of the first week of October (Table 1). Berry skins were separated and analyzed in the same manner as in RNO with the same Illumina technology. Grapes were harvested at a lower °Brix range in BOD (19.5 to 22.5°Brix) than in RNO (20 to 26°Brix) because fruit maturity for making wine is typically reached in the BOD region at a lower sugar level.

Transcript abundance of the RNA-Seq reads from both RNO and BOD was estimated using Salmon software [24] with the assembly and gene model annotation of Cabernet Sauvignon [25, 26]. The TPM (transcripts per million) were computed for each gene from each experimental replicate (n = 3) from berry skins at different sugar levels ranging from 19 to 26°Brix (Additional File 1). Principal component analysis of the transcriptomic data showed clear grouping of experimental replicates with the largest separation by location (principal component 1 (PC1) = 51% variance) and then °Brix (principal component 2 (PC2) = 22% variance) of the berry skin samples (Fig. 1). To get different perspectives of the data, three approaches were used to further analyze the transcriptomic data. One focused on expression at one similar sugar level in both locations. Another identified a common set of genes whose transcript abundance changed in both locations. And the third one was a more comprehensive network analysis using all of the sugar levels and the two locations.

**Figure 1.**
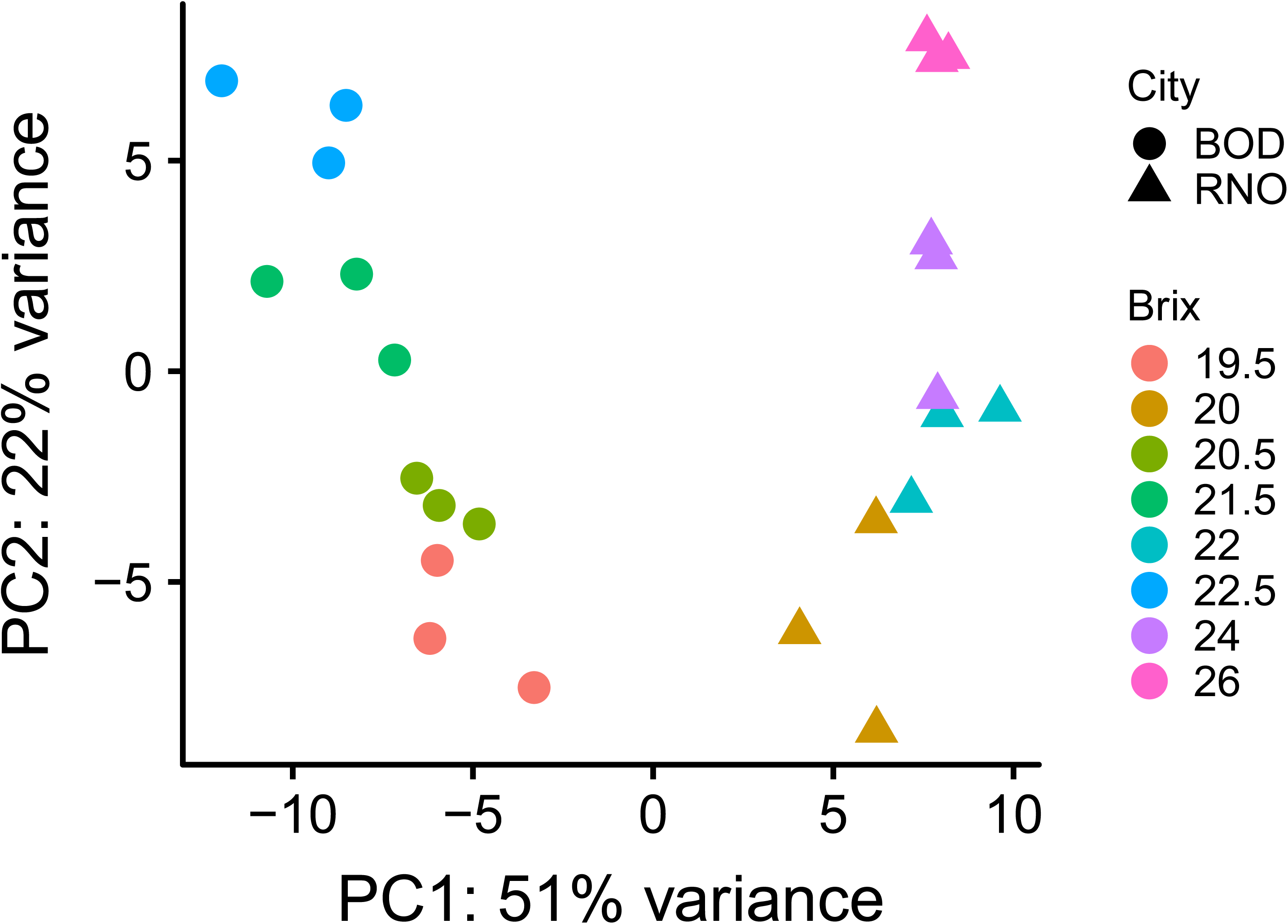
Principal component analysis of the transcriptomic data from berry skin samples at different °Brix levels from BOD and RNO.

Since sugar levels were not exactly the same between the two locations, 5528 differentially expressed genes (DEGs, at an FDR padj-value < 0.05) were identified between the two locations in approach 1 at the sugar level closest to the 22°Brix level (21.5°Brix in BOD vs 22°Brix in RNO) using DESeq2 [27] (Additional file 2). DEGs will refer to this set of genes throughout this manuscript. Gene set enrichment analysis with topGO was determined for these 5528 genes (Additional file 3) and the top gene ontology (GO) categories for biological processes based on the number of genes identified were cellular metabolic process (3126 genes, padj-value = 2.3E-03), biosynthetic process (2371 genes, padj-value = 7.7E-09), and response to stimulus (2324 genes, padj-value = 1.21E-26). Other important and highly significant categories were response to stress (1514 genes, padj-value = 5.69E-24) and developmental process (1280 genes, corrected p-value = 8.09E-12). There were 910 GO categories in total that were significantly enriched (Additional file 3). The relationship between the top 25 GO categories can be seen in Additional file 4. We use the term “significantly” throughout this text to mean statistically significant at or below a padj-value of 0.05. Amongst the top stimulus subcategories with the largest number of genes were response to abiotic stimulus (950 genes; padj-value = 9.1E-29), response to endogenous stimulus (835 genes, padj-value = 1.43E-21; 256 of which were related to response to abscisic acid), response to external stimulus (719 genes, padj-value = 1.08E-24), and biotic stimulus (520 genes, padj-value = 5.29E-22). There were many other environmental stimuli categories with significant gene set enrichment including response to light stimulus (234 genes), response to osmotic stress (171 genes), and response to temperature stimulus (158 genes).

In approach 2, we examined which gene expression was changing with °Brix level in both locations to identify a common set of genes differentially expressed during berry development with very different environmental conditions. The significant differences in transcript abundance in each location was determined with DESeq2 using the lowest °Brix sampling as the control. For example, the control sample in RNO was the lowest sugar sampling at 20 °Brix; the transcript abundance of the three higher °Brix samplings were compared to the transcript abundance of the control. The genes that had significantly different transcript abundance relative to control in at least one of the comparisons were identified in RNO and BOD. These gene lists were compared and the common gene set consisting of 1985 genes for both locations was determined (ap2 tab in Additional file 5). Comparing this common gene list (ap2) to the DEGs from approach 1 identified 907 genes that were common to both sets, indicating that this subset was differentially expressed between the locations at 22°Brix. The other 1078 genes did not differ significantly between locations. This 1078 gene subset list can be found in Additional file 5 (ap2-ap1 tab). The GO categories most enriched in this gene set included response to inorganic substance, response to abiotic stimulus and drug metabolic process.

In approach 3, using a more powerful approach to finely distinguish the expression data for all sugar levels, Weighted Gene Coexpression Network Analysis (WGCNA) identified gene sets common to (based upon correlation) and different gene expression profiles between BOD and RNO. All expressed genes for all °Brix levels (Additional file 1) were used in this analysis. Additional details of the analysis are described in the Materials and Methods section. Twenty-one modules or gene subnetworks were defined (Additional file 6) and a heat map was generated displaying the module-trait relationships (Additional file 7). The grey module is not a real module but a place to put all genes not fitting into a real module; thus, it was not counted as one of the twenty-one gene modules above. Eight modules had similar gene expression profiles for BOD and RNO (padj-value > 0.05); these included cyan, midnightblue, pink, green yellow, salmon, blue, grey60, and royalblue. This gene set consisted of 8017 genes (see ap3 tab in Additional file 5 for the gene list). Comparing this common gene set from the WGCNA with the DEGs from approach 1 revealed that 524 genes in common were found in both sets. This subset was removed from the WGCNA to produce a gene list of 7492 common to both locations and not differing in their transcript abundance at 22°Brix (ap2-ap1). This represents 25% of the total 29,929 genes in all of the modules. This gene set was compared with the ap2-ap1 gene set from approach 1 and 845 genes were found in common in both sets. The remainder from ap2-ap1 provided an additional 232 genes to the common set of genes from ap3-ap1 not affected by location giving a total number of 7724 genes, representing 25.8% of the genes expressed. This gene set is listed in ap2-ap1_union_ap3-ap1 tab in Additional file 5. The GO categories most enriched in this gene set included general categories such as organic substance biosynthetic process and organelle organization. There were 785 enriched GO categories in total.

In approach 3, further analysis of the gene modules using gene set enrichment analysis was performed with genes that had a kME > 0.80 for each module in the WGCNA (Additional file 8). The similar gene sets in common with both locations with decreasing transcript abundance as sugar levels increased (negative correlation with °Brix) were enriched with the GO categories involving growth and water transport (blue module), and translation (grey60 module). The common gene sets with increasing transcript abundance as sugar levels increased were enriched with the GO categories involving gene silencing (cyan module), aromatic compound metabolism (midnight blue), organic substance catabolism (pink module), and DNA metabolism (salmon).

Most modules were positively or negatively correlated with BOD and RNO berries (e.g. black, yellow, red, turquoise, etc.). The turquoise module was the largest module and consisted of 5029 genes; it had the most positive and negative correlations for BOD and RNO, respectively (Additional file 7). This gene set was similar to the DEGs defined by DESeq2 with the largest differences between BOD and RNO at 22°Brix. Gene set enrichment analysis of genes within the turquoise module having a kME of 0.80 or higher (1090 genes) revealed many common GO categories with the DEGs (Additional file 8); 81% (481 of 594) of the GO categories from the turquoise module subset were also found in the 910 GO categories of the DEGs (53% of total). Some of the most enriched GO categories in the turquoise module were organic acid metabolism, flavonoid metabolism, lipid biosynthesis, response to abiotic stimulus, isoprenoid metabolism, response to light stimulus and photosynthesis. The gene expression profiles of this module declined in transcript abundance with increasing sugar levels (negative correlation with °Brix).

The yellow module was another large module (3008 genes) that was the second most positively correlated with BOD. This module was highly enriched with GO categories involving biosynthesis, defense responses and catabolic processes. The *WRKY75* gene (g104630; this g# term is used as an abbreviated gene loci name in the Cabernet Sauvignon genome throughout this paper) was in the top 4 hub genes (kME = 0.97) in the yellow module (Additional file 6). WRKY75 is a transcription factor that positively regulates leaf senescence. It is induced by ethylene, ROS (reactive oxygen species) and SA (salicylic acid) and is a direct target of EIN3 (ethylene insensitive 3) [28].

The green (2287 genes) and brown (4147 genes) modules were also large modules that were most positively correlated with RNO (0.92 and 0.9, respectively). The green module was highly enriched in the GO category involving response to chemical. The brown module was highly enriched in GO categories involving multiple catabolic processes.

Thus, the WGCNA results, which utilized all of the expressed genes from all °Brix levels were consistent with the DESeq2 results that only compared transcript abundances at 22°Brix or between locations. The WGCNA results were more comprehensive and complemented the results of approaches 1 and 2 by identifying hub genes and gene subnetworks. These subnetworks were linked and further defined by their highly correlated coexpression profiles and enriched GOs.

## Transcriptomic profiles dynamically changing with sugar levels

### DEGs with largest increases in transcript abundance between sugar levels

As a first approach to examining the 5528 DEG dataset, differences between the transcript abundance in berries with the lowest and highest °Brix levels at the two locations were determined (Additional file 2). Eight examples of the many DEGs with the largest transcript abundance differences from BOD and RNO (*EXL2* (exordium like 2, g068700), *HB12* (homeobox 12, g223410), *BSMT1* (benzoate/salicylate methyltransferase 1, g336810), *HAD* (haloacid dehalogenase-like hydrolase protein, g070140), *STS24* (stilbene synthase 24, g435870), *NAC073* (NAC domain containing protein 73, g125400), *TPS35* (terpene synthase 35, g087040), and *MAT3* (methionine adenosyltransferase 3, g013310)) were selected and are presented in Figure 2. The major point of showing this plot is to highlight the general trends of continuously increasing transcript abundance with sugar levels for these genes; half of these genes start at similar transcript abundance levels around 20°Brix for both BOD and RNO berries and increase in transcript abundance at a higher rate in BOD grapes as sugar levels increase. The other half increased at approximately the same rate for both locations but had higher amounts at the BOD location at the same sugar levels. These data were fitted by linear regression to compare the slopes of the lines. The slopes were significantly higher for *EXL2*, *BSMT1*, *STS24*, and *TPS35* for BOD as compared to RNO berries, but not for the other four genes (data not shown). The significantly increased rate of change for transcript abundance in the BOD berries indicated that the berries in BOD may have ripened at a faster rate relative to sugar level.

**Figure 2.**
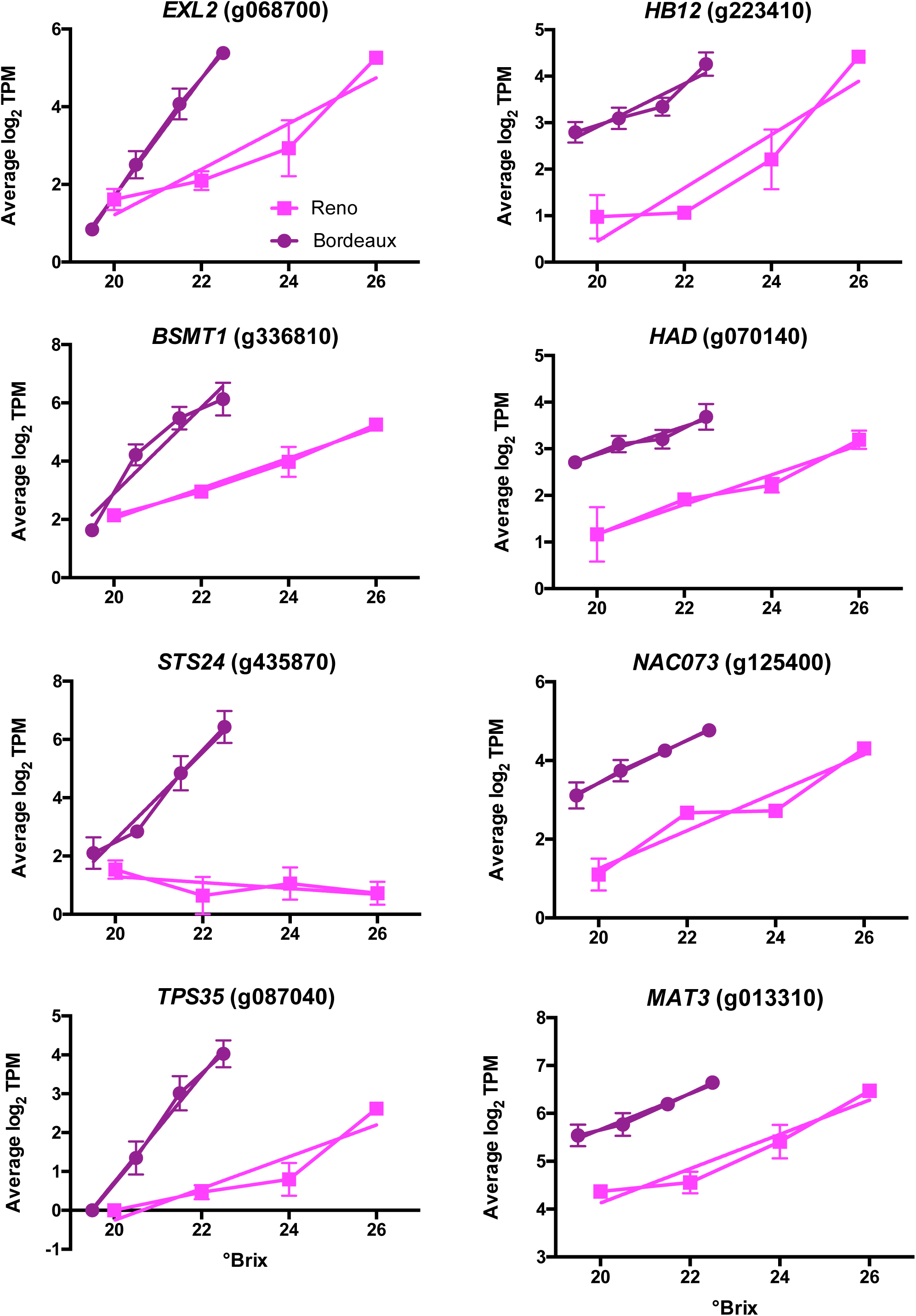
Plots of the top genes from berry skins from Bordeaux and Reno with the highest increase in transcript abundance (TPM) between the lowest and highest sugar levels (°Brix). Values are the means ± SE (n = 3). Error bars not shown are smaller than the symbol. The symbol legend is displayed in the figure. EXL2 is EXordium Like 2; HB12 is HOMEOBOX 12; BSMT1 is a benzoate/salicylate methyltransferase 1; HAD is haloacid dehalogenase-like hydrolase protein; STS24 is stilbene synthase 24, NAC073 is a NAC domain containing protein; TPS 35 is terpene synthase 35, and MAT3 is methionine adenosyltransferase 3.

To get deeper insights into these dynamic gene sets from BOD and RNO, gene set enrichment analysis of the top 400 DEGs with the greatest increase in transcript abundance from the lowest sugar level to the highest sugar level was performed for each location. The top 400 DEGs for BOD berries were highly enriched in biosynthetic processes involving amino acid and phenylpropanoid metabolism (Additional File 9); defense responses, response to fungus and response to ethylene stimulus were other highly enriched categories. The top 400 DEGs for RNO berries were enriched in response to oxygen-containing compound, response to hormone and response to abscisic acid (Additional File 10).

### DEGs with largest decreases in transcript abundance between sugar levels

Eight examples of DEGs with the greatest decrease in transcript abundance with increasing sugar levels are presented in Figure 3. They include lipid and cell wall proteins (e.g. extensin like and lipid transferase proteins) and an aquaporin (TIP1;1, tonoplast intrinsic protein 1;1). The data were fitted to linear regressions and the slopes statistically compared between BOD and RNO berries. In some cases, the slopes of the linear regression lines of the DEGs were not statistically different (bifunctional inhibitor lipid transfer protein and *DUF642* (domain of unknown function 642); but in the other cases presented, there were similar amounts of transcript abundance around 20°Brix, but there were significantly different slopes. Again, there is a trend for the transcript abundance of many of these genes to change more significantly in BOD berries than RNO berries relative to sugar level.

**Figure 3.**
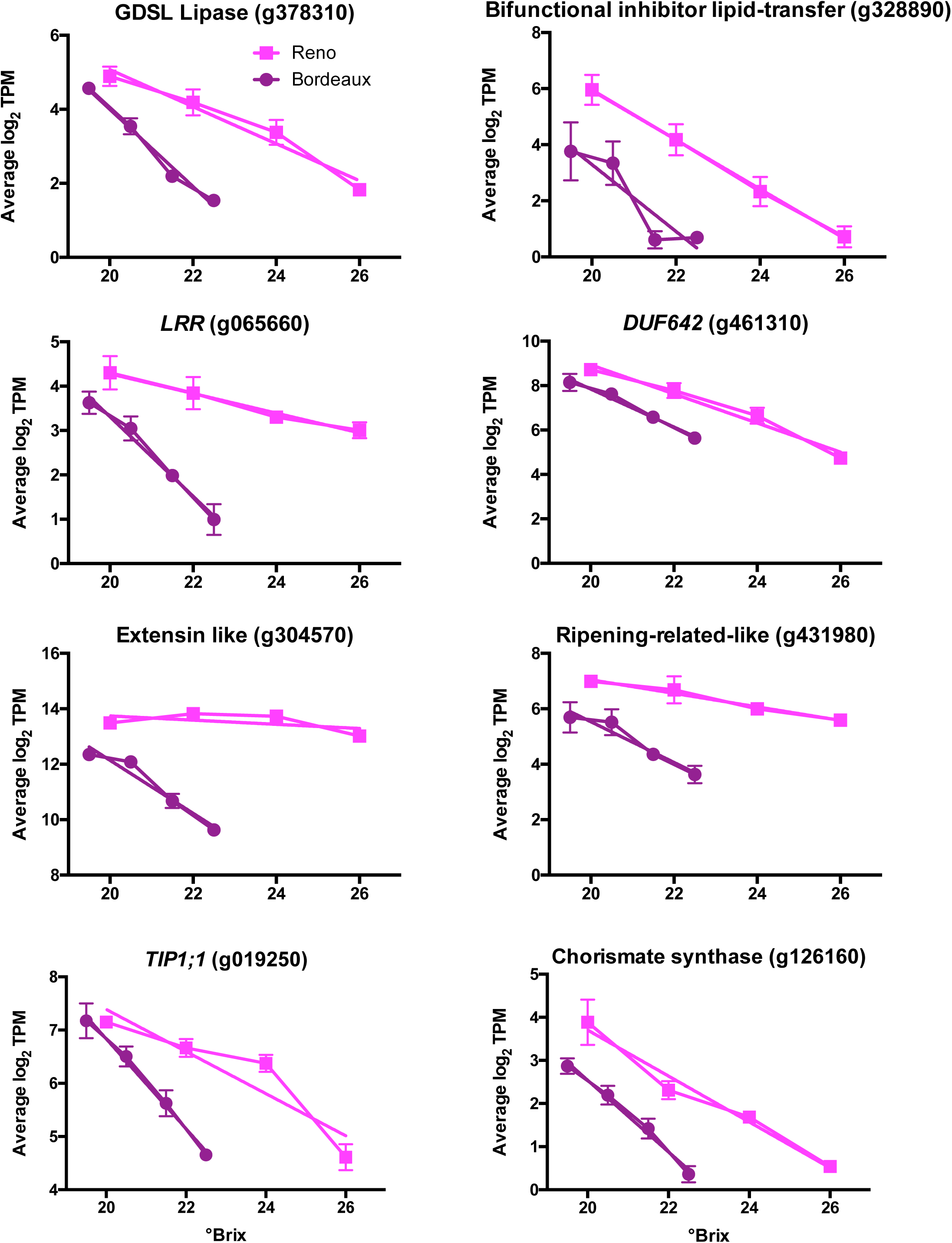
Plots of the top genes from berry skins from Bordeaux and Reno with the highest decrease in transcript abundance (TPM) between the lowest and highest sugar levels (°Brix). Values are the means ± SE (n = 3). Error bars not shown are smaller than the symbol. The symbol legend is displayed in the figure. GDSL is a sequence motif; LRR is a leucine rich repeat protein; DUF642 is a domain of unknown function protein; TIP1;1 is a tonoplast intrinsic protein 1;1.

### Differences in autophagy genes between BOD and RNO

Berry ripening is a degradative process involving autophagy. As a general rule, the transcript abundance of genes involved in autophagy increased with sugar level and had a higher transcript abundance in BOD berries relative to RNO berries at the same sugar level (Fig. 4), consistent with the hypothesis that BOD berries ripened faster than RNO berries relative to sugar level.

**Figure 4.**
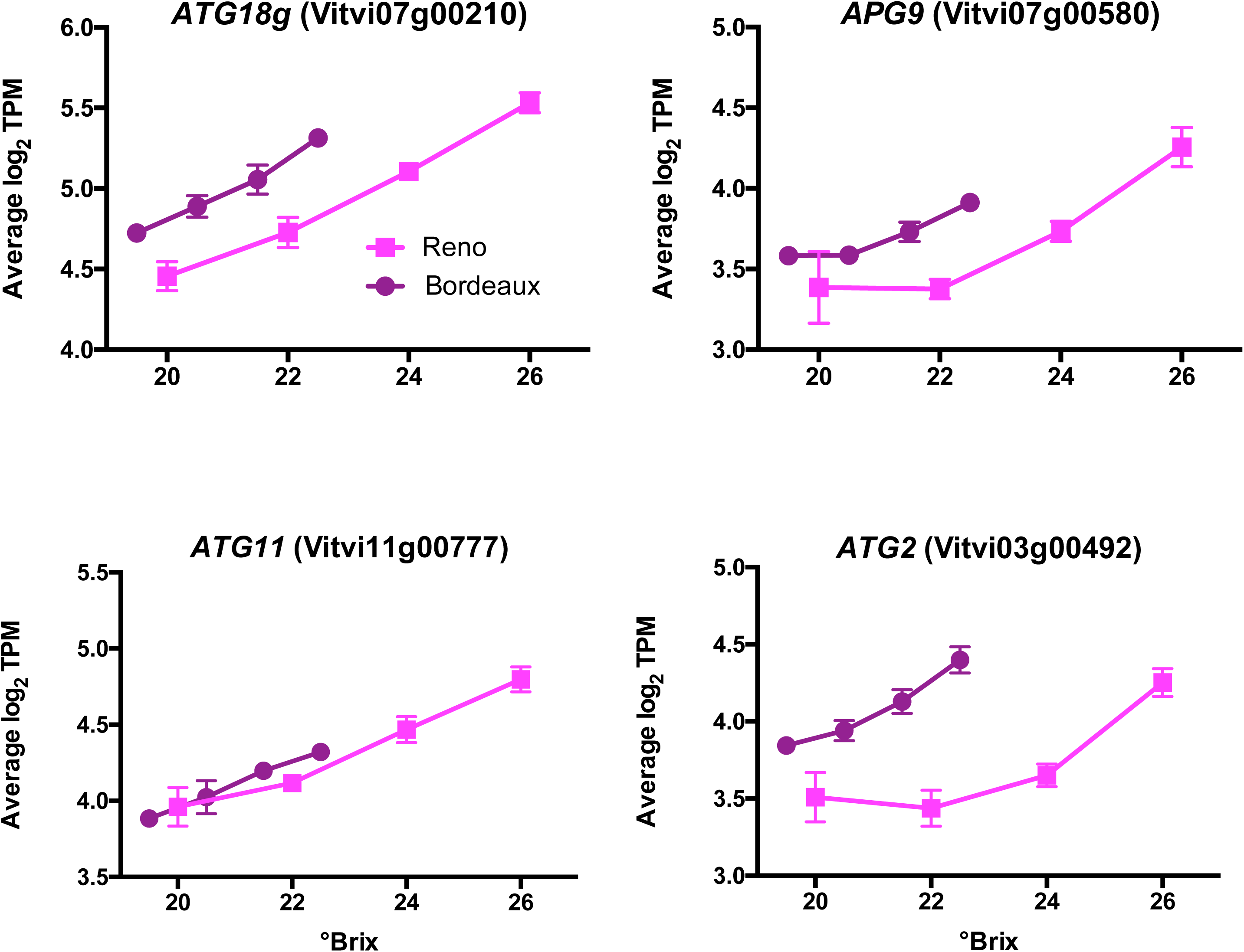
The transcript abundance of some autophagy-related (ATG) genes. Data shown are the means ± SE; n = 3. Error bars not shown are smaller than the symbol. The symbol legend is displayed in the figure. APG9 is Autophagy 9.

## Aroma- and flavor-associated DEGs

Many aroma and flavor-associated compounds are synthesized in the late stages of berry ripening and sensitive to the environment. The major metabolic pathways affecting flavor and aromas in grapes and wines include the terpenoid, carotenoid, amino acid, and phenylpropanoid pathways [6]. These pathways were identified by topGO to be highly enriched in the DEGs and the turquoise module. Some of the DEGs are involved in these pathways and will be presented in the following subsections.

### Terpene synthase genes with the greatest transcript abundance differences between BOD and RNO

Cultivar differences in berries are often ascribed to differences in aroma compounds. One of the main classes of cultivar specific aroma compounds is the terpene group [29]. The transcript abundance of a number of terpene synthases were higher in BOD berries as compared to RNO berries (Fig. 5). All but one of these (terpene synthase 55; *TPS55*) increased in transcript abundance with increasing sugar levels. TPS35 is a β-ocimine synthase (β-ocimine is a main component of snapdragon flower aroma [30]). TPS08 is a γ-cadinene synthase, TPS26 is a cubebol/δ-cadinene synthase, TPS4 and 10 are (E)-α-bergamotene synthases, and TPS07 and 28 are germacrene-D synthases; these enzymes produce sesquiterpenes found in essential plant oils (see [29] and references therein for the function of all of these terpenoid genes). TPS55 is a linalool/nerolidol synthase which synthesizes acyclic terpene alcohols; linalool contributes significantly to the floral aromas of grape berries and wines. TPS68 is a copalyl diphosphate synthase involved in diterpenoid biosynthesis and TPS69 is an ent-kaurene synthase. Both TPS68 and 69 are diterpene synthases and are part of the ent-kaurene biosynthesis pathway.

**Figure 5.**
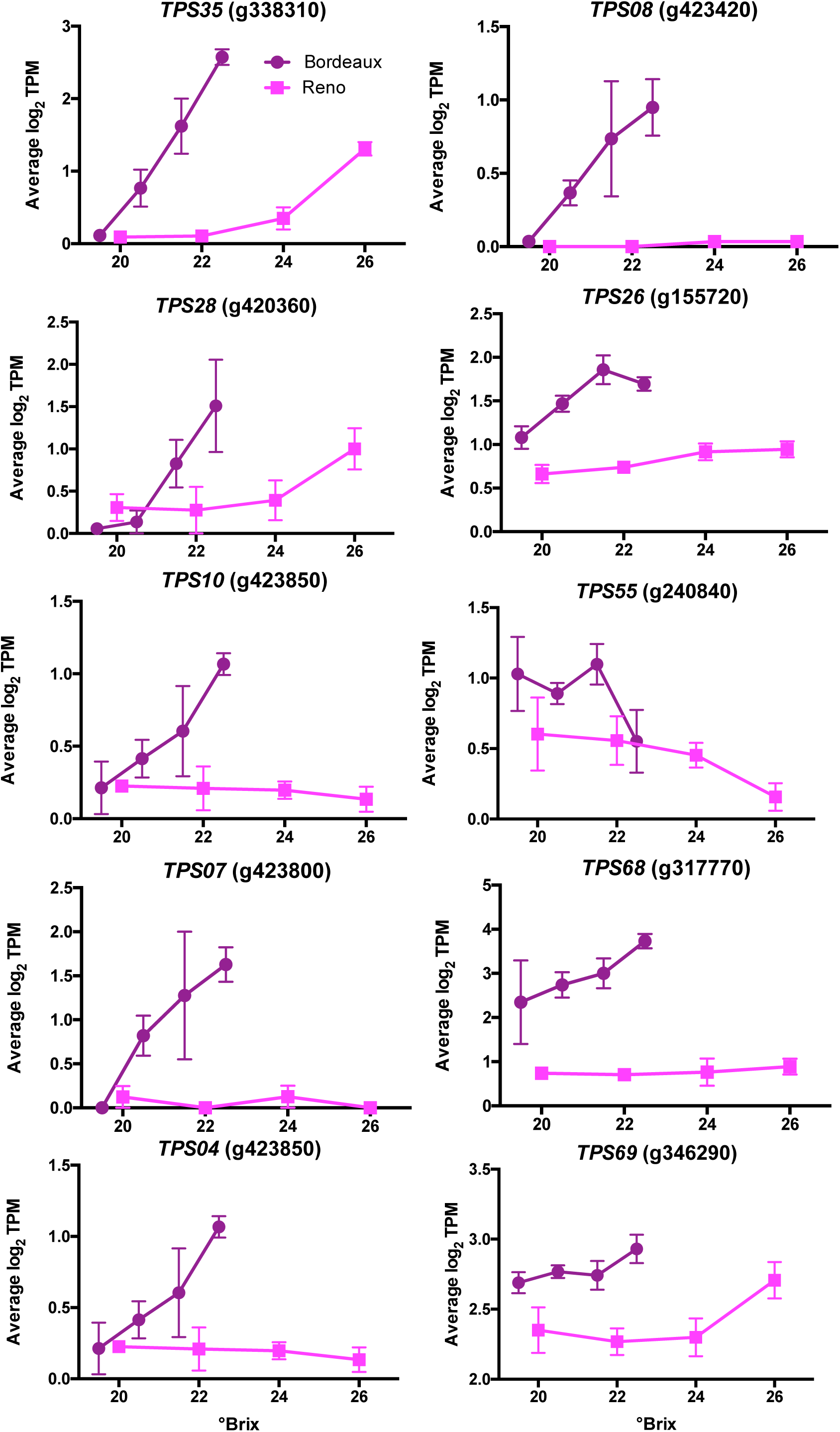
Expression profiles of some terpene synthase (TPS) genes. Data shown are the means ± SE; n = 3. Error bars not shown are smaller than the symbol. The symbol legend is displayed in the figure.

### Other terpenoid and carotenoid metabolism-related genes

Carotenoid metabolism is another biosynthetic pathway that contributes to flavor and aroma in grapes [31]. There are a number of key genes that contribute to terpenoid and carotenoid metabolism that have a higher transcript abundance (Additional file 2) in BOD berries as compared to RNO berries. For example, DXR (1-deoxy-D-xylulose 5-phosphate reductoisomerase, g360850) catalyzes the first committed step and HDS (4-hydroxy-3-methylbut-2-en-1-yl diphosphate synthase; g379980) enzyme controls the penultimate steps of the biosynthesis of isopentenyl diphosphate (IPP) and dimethylallyl diphosphate (DMAPP) via the methylerythritol 4-phosphate (MEP) pathway. Other examples are two phytoene synthases (PSY), g180070 and g493850); PSY is the rate-limiting enzyme in the carotenoid biosynthetic pathway.

### Amino acid and phenylpropanoid metabolism genes

Amino acids contribute to the aroma and flavor of grapes and wines [7]. The amino acid metabolism functional GO category is highly enriched in the group of DEGs between BOD and RNO (Additional file 3) and more specifically in the top 400 BOD DEGs (Additional file 9). Some examples of genes involved in amino acid metabolism that have a higher transcript abundance in BOD berries (see Fig. 6) are phenylalanine ammonia lyase 1 (*PAL1*, g533070 and eight other paralogs can be found in Additional file 2), which catalyzes the first step in phenylpropanoid biosynthesis, branched-chain-amino-acid aminotransferase 5 (*BCAT5*, g220210), which is involved in isoleucine, leucine and valine biosynthesis, 3-deoxy-D-arabino-heptulosonate 7-phosphate synthase 1 (*DHS1*, g082490), which catalyzes the first committed step in aromatic amino acid biosynthesis), and tyrosine aminotransferase 7 (*TAT7*; g116950), which is involved in tyrosine and phenylalanine metabolism. Included in this group were 44 stilbene synthases (*STS*), which are part of the phenylpropanoid pathway; these *STSs* had a higher transcript abundance in BOD berries as compared to RNO berries, with very similar transcript abundance profiles to *PAL1* (see Additional file 11 for two typical examples).

**Figure 6.**
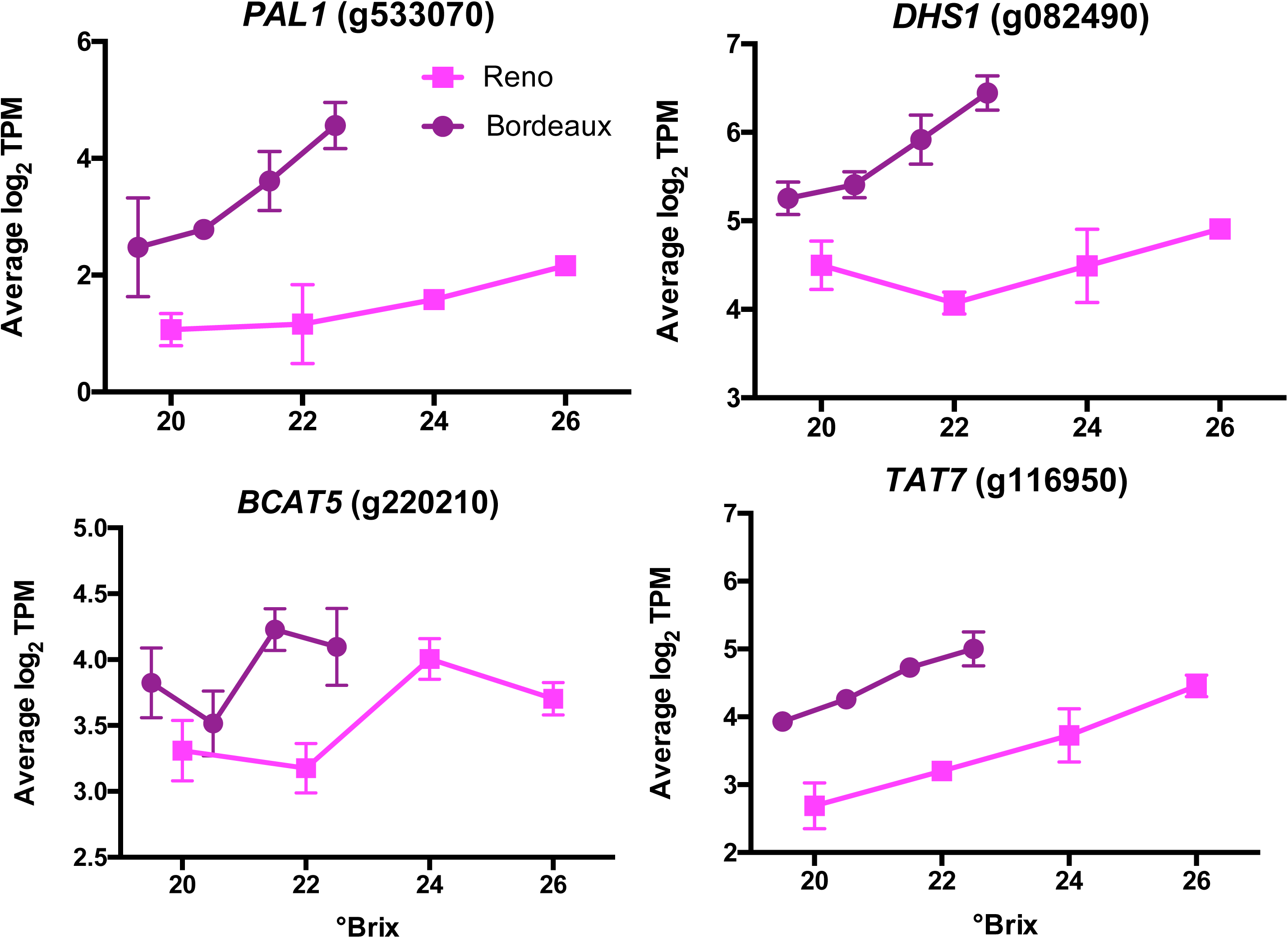
The transcript abundance of some DEGs that are involved in the amino acid metabolism. Data shown are the means ± SE; n = 3. Error bars not shown are smaller than the symbol. The symbol legend is displayed in the figure. PAL1 is a phenylalanine ammonia lyase; DHS1 is a 3-deoxy-D-arabino-heptulosonate 7-phosphate synthase; BCAT5 is a branched-chain-amino-acid aminotransferase; and TAT7 is a tyrosine aminotransferase.

## DEGs associated with abiotic stimuli

### Light-responsive genes

In a previous analysis, WGCNA defined a circadian clock subnetwork that was highly connected to transcript abundance profiles in late ripening grapevine berries [4]. To compare the response of the circadian clock in the two different locations, we plotted all of the genes of the model made earlier [4]. Most core clock genes (Additional file 12) and light sensing and peripheral clock genes (Additional file 13) had significantly different transcript abundance in BOD berries than that in RNO berries at the same sugar level (profiles bracketed in a red box). All but one of these (*PHYC*, phytochrome C, g088040) had higher transcript abundance in BOD berries relative to RNO berries. The transcript abundance of other genes had nearly identical profiles (not bracketed in a red box). These data are summarized in a simplified clock model (Fig. 7), which integrates PHYB as a key photoreceptor and temperature sensor [32, 33] that can regulate the entrainment and rhythmicity of the core circadian clock, although to be clear it is the protein activity of PHYB, not the transcript abundance that is regulating the clock.

**Figure 7.**
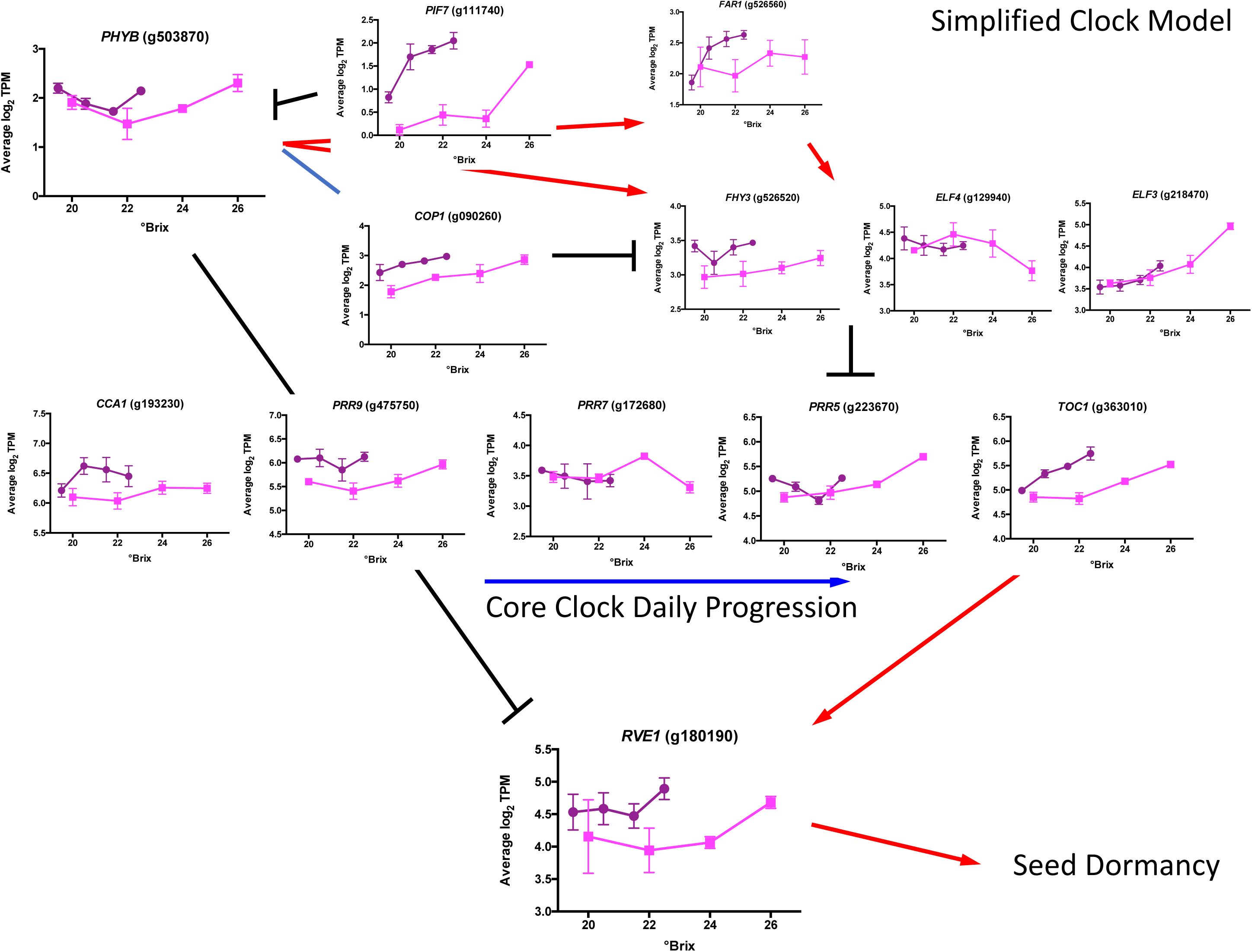
A simplified model of the core circadian clock genes. Black lines and bars represent known inhibitory reactions, red arrows indicate known stimulatory reactions, and blue lines represent known physical interactions. PHYB is phytochrome B; PIF7 is Phytochrome Interacting Factor 7; COP1 is Constitutive Photomorphogenic 1; FAR1 is Far-Red Impaired Response 1; FHY3 is Far Red Elongated Hypocotyl 3; ELF4 is Early Flowering 4; ELF3 is Early Flowering 3; CCA1 is Circadian Clock Associated 1; PRR9 is Psuedo-Response Related 9; PRR7 is Psuedo-Response Related 7; PRR5 is Psuedo-Response Related 5; TOC1 is Timing of CAB expression 1; and RVE1 is Reveille 1.

### Chilling-responsive genes

Temperatures were colder in RNO than BOD, reaching chilling temperatures in the early morning hours. A number of previously identified chilling-responsive genes [34] in Cabernet Sauvignon had a higher transcript abundance in RNO berries as compared to BOD berries (Fig. 8). These genes included *CBF1* (C-repeat/DRE binding factor 1, g435450; previously named *CBF4*, but renamed to be consistent with the ortholog of Arabidopsis), a transcription factor that regulates the cold stress regulon [35], *IDD14* (Indeterminate-Domain 14, g000790), a transcription factor that generates an inhibitor to regulate starch metabolism [36], *CML41* (calmodulin-like 41, g041290), that encodes a calmodulin-like protein, *CYSB* (cystatin B, g023260), a cysteine proteinase inhibitor that confers cold tolerance when overexpressed [37], *XTH23* (xyloglucan endotransglucosylase/hydrolase 23, g572510), that encodes a cell wall loosening enzyme, and *SULTR3;4* (sulfate transporter 3;4, g392710).

**Figure 8.**
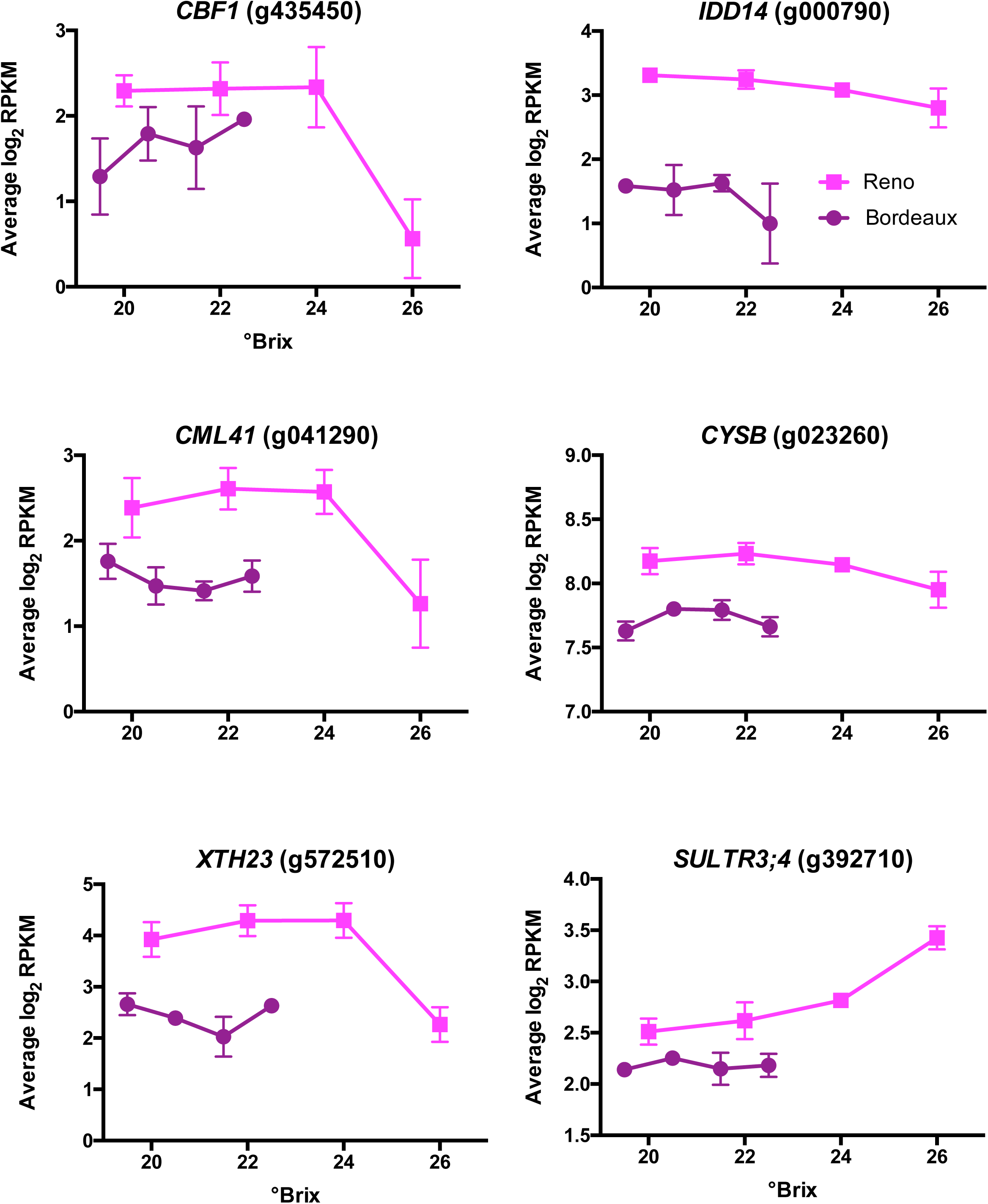
Transcript profiles of some cold responsive genes. Data shown are the means ± SE; n = 3. Error bars not shown are smaller than the symbol. The symbol legend is displayed in the figure. CBF1 is C-Repeat Binding Factor 1; IDD14 is Indeterminate-Domain 14; CML41 is Calmodulin 41; CYSB is Cystatin B; XTH23 is Xyloglucan Endotransglucosylase/Hydrolase 23 and SUFTR3;4 is Sulfate Transporter 3;4.

## DEGs associated with biotic stimuli

The top DEGs in BOD berries were highly enriched in biotic stimuli genes. Some of these genes included pathogenesis proteins (*PR*); these genes had higher transcript abundance in BOD berry skins relative to RNO berries (Fig. 9). The transcript abundance of *PR10* (g212910) increased with the sugar level. The transcript abundance of this *PR10* gene increases in response to powdery mildew (*Erysiphe necator*) inoculation in Cabernet Sauvignon leaves [38]. Furthermore, many other genes induced by powdery mildew were also at much higher transcript abundance levels in BOD berries. These included a PR3 protein that is a class IV chitinase, a thaumatin-like protein (PR5) and a number of the stilbene synthases mentioned in the phenylpropanoid metabolism section above (see Additional file 11). *MLA10* (Intracellular Mildew A 10, g343420; Affymetrix probe set 1615715_at in [38]) matches to a fungal protein from *E. necator* and was used as a control probe set for the presence of powdery mildew [38]. g343420 had a much higher transcript abundance in BOD berries than that in RNO berries. These data indicate that powdery mildew infection may have been higher in BOD berries and that this infection may induce some of the phenylpropanoid pathway as well.

**Figure 9.**
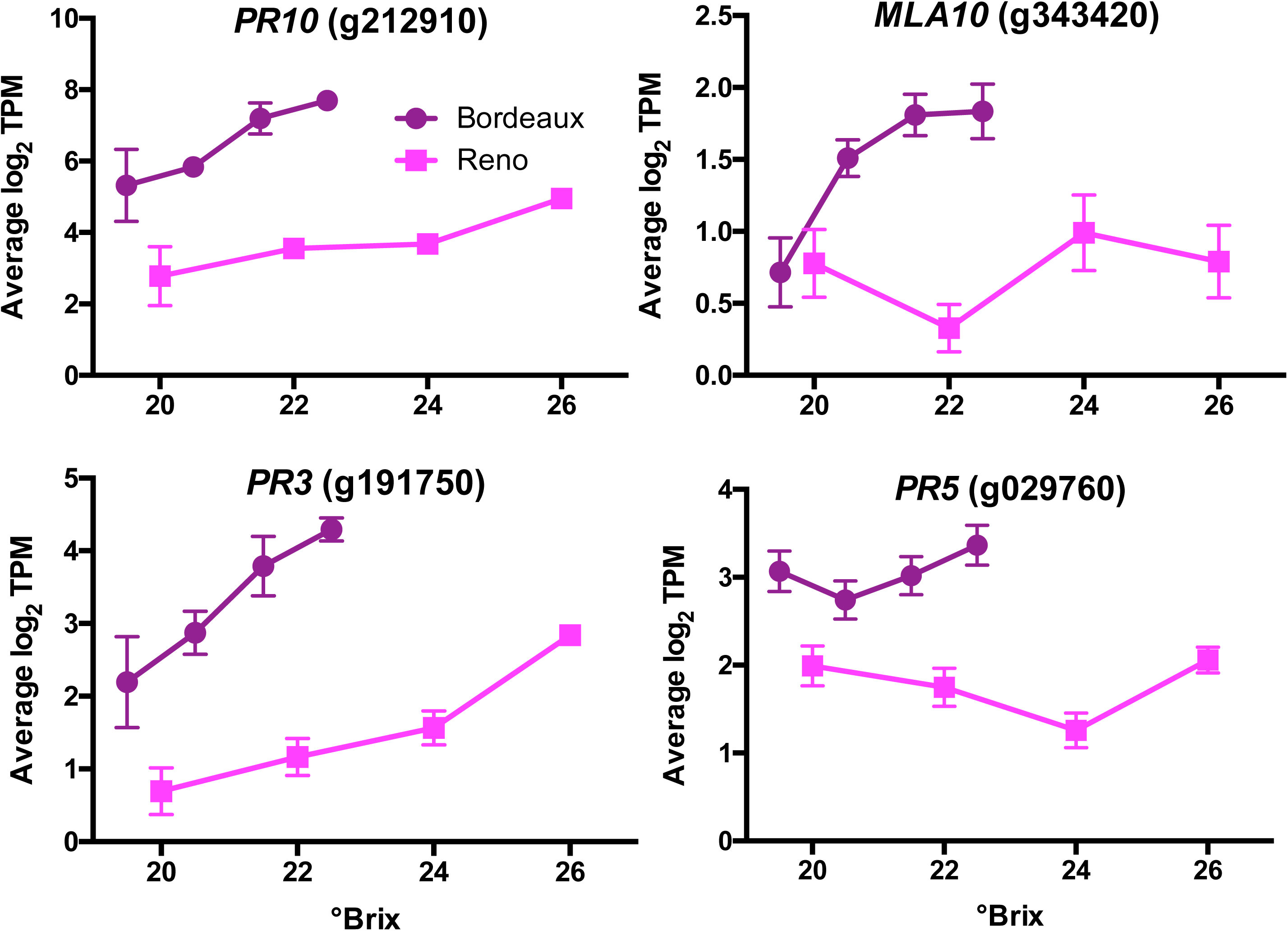
Expression profiles of pathogenesis proteins (PR) involved with powdery mildew. Data shown are the means ± SE; n = 3. Error bars not shown are smaller than the symbol. The symbol legend is displayed in the figure. MLA10 is Intracellular Mildew A 10.

## DEGs associated with hormonal stimuli

### Auxin signaling genes

Auxin transport (38 genes; padj-value = 4.43E-08) and cellular response to auxin stimulus (45 genes, padj-value = 9.12E-05) were highly enriched GO categories for the DEGs (Additional file 3). Auxin is known to have multiple effects on grape berry ripening [39, 40]. Auxin can delay berry ripening at the veraison stage, which is at the beginning of berry ripening. Some auxin metabolism (*GH3.1*; GH3 family protein, g538930) and signaling genes such as *IAA13* (indole acetic acid 13, g527400), *IAA27* (g326620), and *ARF5* (auxin-response factor, g075570) had a higher transcript abundance in RNO berries (Additional file 1). Other auxin metabolism (*GH3*.6, *JAR1* (jasmonate resistant 1), g170030) and auxin signaling genes had a higher transcript abundance in BOD including *ARF2* (g469780), *ARF8* (g180460), *ARF11* (g380160), *IAA16* (g318830), *ARAC1* (Arabidopsis RAC-like 1, g320970), and *GID1B* (gibberellic acid insensitive dwarf1B, g071190), a gibberellin receptor (Additional file 1).

### ABA metabolism and signaling genes

ABA is a stress hormone that responds to water deficits in grapevine [41]. A number of ABA-related genes are differentially expressed in berry skins between the two locations (Additional file 14). *NCED3* (nine-cis epoxycarotenoid dioxygenase 3, g221190) and *NCED5* (g404590), which are responsive to water deficit [42, 43], had higher transcript abundance in RNO and *NCED6* (g203160), which is highly expressed in embryos [42], had higher transcript abundance in BOD. NCED6, but not NCED3 is involved in seed ABA and seed dormancy [43]. Additionally a number of other genes involved in the ABA signaling pathway had higher transcript abundance in BOD including *ABF2* (abscisic acid responsive elements binding factor 2, g286950), *ABF4* (g312300) and *ABCG40* (adenosine triphosphate binding casette G 40, g143240) [44, 45]. Interestingly, BAM1 (Barley Any Meristem 1) was identified to be the receptor to a root signaling peptide hormone (CLE25, clavata3/esr-related 25) that responds to water deficit and upregulates *NCED3* transcript abundance in Arabidopsis leaves [46]. The transcript abundance of *BAM1* (g220020) was significantly higher in RNO berries than that of BOD berries (Additional File 14). There were no significant differences in the transcript abundance of *CLE25* (g007470); it was highly variable.

### Ethylene signaling genes higher in BOD

There were 71 DEGs that were enriched in the response to ethylene GO category (Additional file 3). Ethylene is a stress hormone that responds to many types of biotic [47] and abiotic [48] stresses in addition to its role in fruit development and ripening [49]. Many ethylene-related genes had a higher transcript abundance in BOD berries. These included ethylene biosynthesis, ethylene receptors and ERF (ethylene response factor) transcription factors (Fig. 10). ERF1 and ERF2 are at the beginning of the ethylene signaling pathway and are direct targets of EIN3 [45, 50]. Other ERF transcription factors (e.g. *ERF98*; g156210) identified as hubs in the ethylene signaling pathway in Arabidopsis leaves [51] were also differentially expressed in a similar manner as *ERF1* (g060690) and *ERF2* (g482650) between the two locations (data not shown).

**Figure 10.**
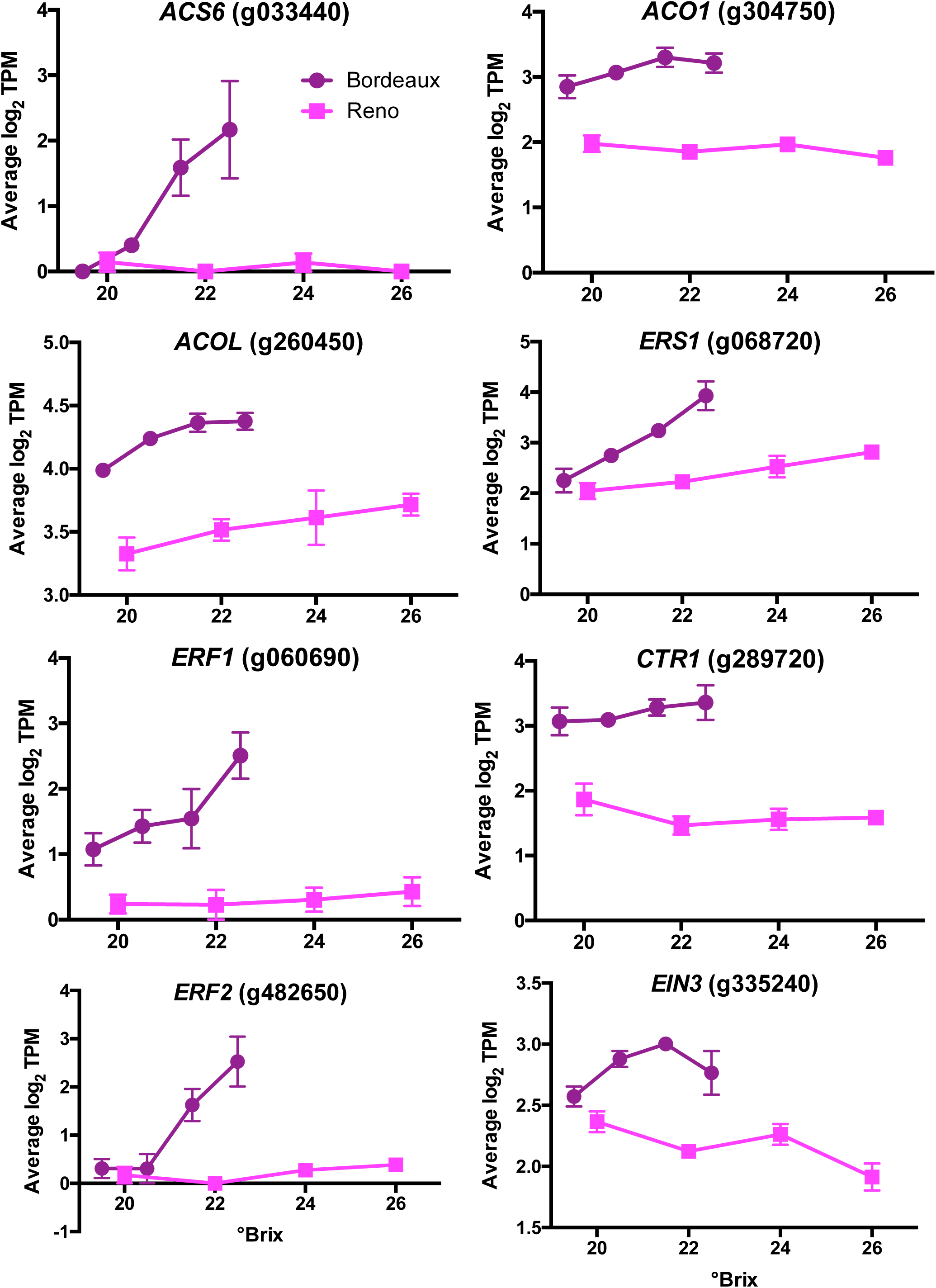
The transcript abundance of DEGs involved with ethylene metabolism and signaling. Data shown are the means ± SE; n = 3. Error bars not shown are smaller than the symbol. The symbol legend is displayed in the figure. ACS6 is 1-Aminocyclopropane-1-Carboxylic Acid (ACC) Synthase 6; ACO1 is ACc Oxidase 1; ACOL is ACc Oxidase-Like; ERS1 is Ethylene Response Sensor 1; CTR1 is Constitutive Triple Response 1, EIN3 is Ethylene-Insenstive 3; and ERF1 and ERF2 are Ethylene Response Factors 1 and 2.

## DEGs associated with mineral nutrients

### Iron-related genes

Fourteen DEGs were associated with genes enriched in response to iron ion (Additional file 3); Eight examples of DEGs involved in iron homeostasis are shown in Figure 11. Iron homeostasis genes *SIA1* (salt-induced ABC kinase, g336700), *VIT1* (vacuolar iron transporter 1, g001160), *ATH13* (Arabidopsis thaliana ABC2 homolog 13, g146610)*, IREG3* (iron regulated 3, g098530), *and ABCI8* (ATP-binding cassette I8, g163790) have higher transcript abundance in BOD berries than in RNO berries. Iron homeostasis genes *YSL3* (yellow stripe-like 3, g223320), *FER1* (ferritin 1, g606560), and *NRAMP3* (natural resistance-associated macrophage protein 3, g413920) had higher transcript abundance in RNO berries compared to BOD berries. Several other ferritin genes were expressed similarly to *FER1* (data not shown). Average available iron soil concentrations were about 5 times higher in the BOD vineyard soil compared to the RNO vineyard soil (Table 1).

**Figure 11.**
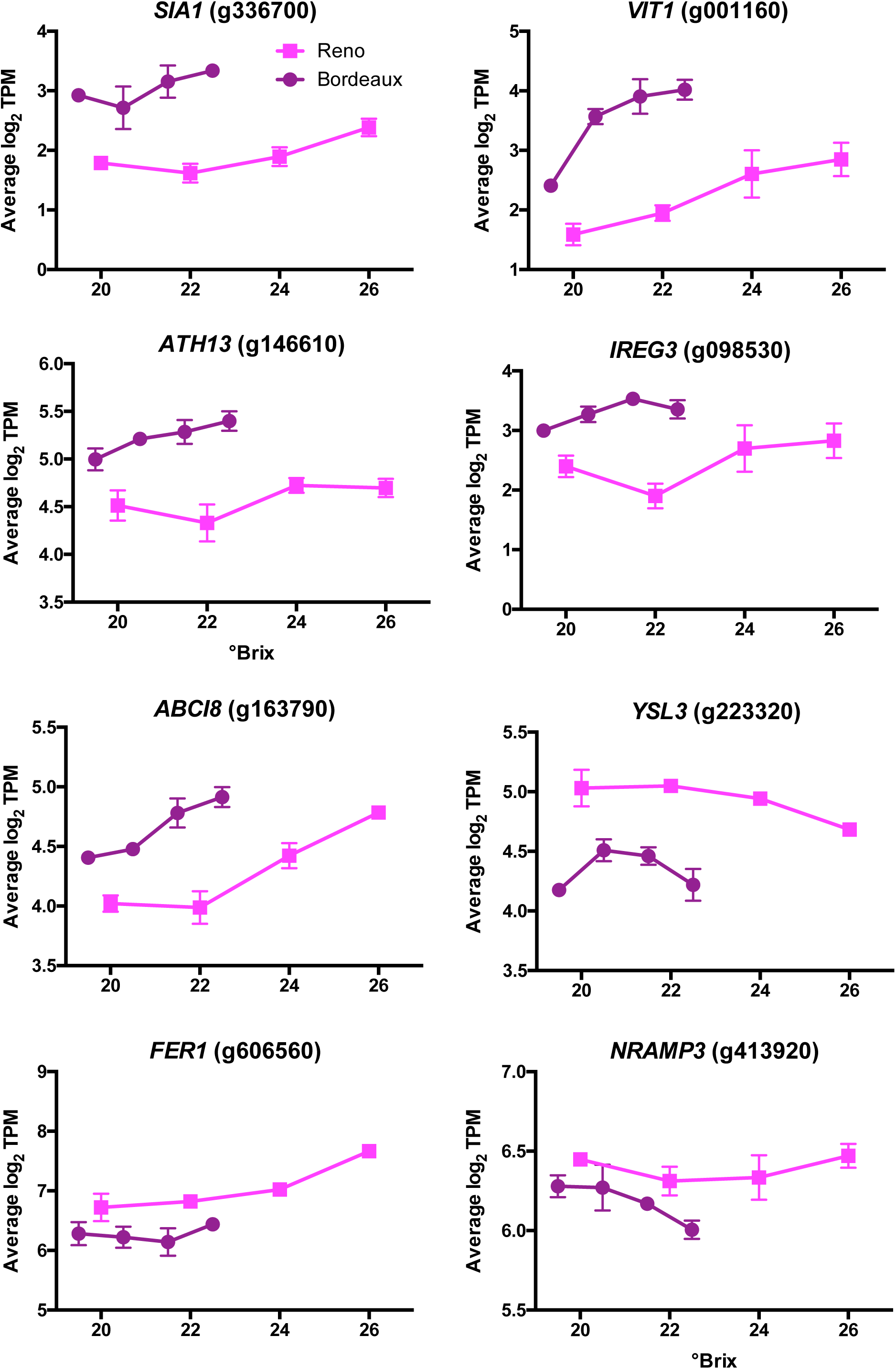
Expression profiles of DEGs involved in iron homeostasis. Data shown are the means ± SE; n = 3. Error bars not shown are smaller than the symbol. The symbol legend is displayed in the figure. SIA1 is Salt-Induced ABC1 Kinase 1; VIT1 is Vacuolar Iron Transporter 1; FER1 and 2 are Ferritin 1 and 2; YSL3 is Yellow Stripe-like 3; IREG3 is Iron-Regulated Protein 3; NRAMP3 is Natural Resistance-Associated Macrophage Protein 3; ATH13 is ABC2 homolog 13; and ABCI8 is ATP-Binding Cassette I8.

## Discussion

The common gene set (ap2-ap1_union_ap3-ap1) for both locations represented approximately 25% of the genes differentially expressed with sugar level or location. Presumably these gene sets represent genes that were not influenced by location (environment) but were influenced by berry development or sugar level. As more locations are compared in the future, these gene sets will likely be reduced in size even further. The processes involved in these gene sets or modules included the increase of catabolism and the decline of translation and photosynthesis. It is clear that these processes play important roles in berry ripening. Most of the genes in the genome varied in transcript abundance with increasing sugar levels and berry maturation and most of these varied with the vineyard site. Many of the DEGs were enriched with gene ontologies associated with environmental or hormonal stimuli.

### DEG expression profiles of grape berry skins were associated with environmental factors and seed development

Plants are exposed to a multitude of factors that influence their physiology even in controlled agricultural fields such as vineyards. The vineyards in BOD and RNO are exposed to very different environments; these environmental influences were reflected in some of the DEG sets with enriched gene ontologies. Our data are consistent with the hypothesis that transcriptomic dynamics in the late stages of berry ripening are sensitive to local environmental influences on the grapevine.

While most transcript abundances in berries are largely influenced by genetics or genotype, environment also plays a large role [2]. It is impossible with the experimental design of this study to determine the amount that each of the environmental factors contributed to the amount of differential expression in these two locations. There were too many variables and too many potential interactions to determine anything conclusively. All we can say is that these genes were differentially expressed between the two locations, which were likely due to known and unknown factors (a sense of place or terroir).

In this study we compared two different clones of Cabernet Sauvignon, one on rootstock and the other on its own roots. There were likely to be some effects on the transcript abundance in the berries between these grapevines as a result of the genetic differences between their roots and their clonal shoots/scions. Not knowing what these genes might be from previous studies prevents us from drawing any clues. These and other factors most certainly affected the berries to some degree. However, our data indicated that grape berry skins also were responsive to a multitude of potential environmental factors in the two vineyard locations and possibly from signals coming from the maturing seed. We say potential environmental factors because we did not control for these factors; we associated transcript abundance with the factors that were different in the two locations. The transcript abundance profiles along with functional annotation of the genes gave us clues to factors that were influencing the berries and then associations were made with the known environmental variables. Further experiments are required to follow up on these observations.

The environmental factors that we were able to associate with differences in transcript abundance (DEGs) between the two locations included air temperatures, light, moisture (rainfall and relative humidity) and biotic stress. These factors in turn were associated with transcript abundance involved with physiological responses and berry traits such as seed and embryo development, hormone signaling (ABA, ethylene and auxin), phenylpropanoid metabolism, and the circadian clock. In the following sections we discuss in more detail some of the possible environmental factors that were reflected in the enriched gene ontologies found in the gene sets from this study.

### Transcript abundance of light-responsive genes

Light regulates the transcript abundance of many genes in plants. It has been estimated that 20% of the plant transcriptome is regulated by white light and this includes genes from most metabolic pathways [52]. Light is sensed by a variety of photoreceptors in plants [33]; there are red/far red, blue and UV light receptors. PHYB is a key light sensor, regulating most of the light sensitive genes [53] and sensing the environment through red light to far-red light ratios and temperature [33, 54]. PHYB entrains the circadian clock affecting the rate of the daily cycle [55] and the expression of many the circadian clock genes [53]; PHYB induces morning phase genes and represses evening phase genes. Other photoreceptors can entrain the circadian clock as well [33].

PHYB and the circadian clock are central regulators of many aspects of plant development including seed germination, seedling growth, and flowering [33, 55, 56]. The circadian clock influences the daily transcript abundance of genes involved in photosynthesis, sugar transport and metabolism, biotic and abiotic stress, even iron homeostasis [55].

Light signaling was very dynamic in the berry skin transcriptome in the late stages of berry ripening with a higher transcript abundance of many light signaling genes in BOD berries. Many photoreceptors that interact with the circadian clock had a higher gene expression in BOD berries. In the circadian clock model, Circadian Clock Associated 1 (*CCA1*) is an early morning gene and has its highest expression at the beginning of the day. It is at the start of the circadian core clock progression through the day, whereas the transcript abundance of Timing Of CAB Expression 1 (*TOC1*) is highest at the end of the day and finishes the core clock progression (Fig. 7). In both of these cases, there is a higher transcript abundance of these genes in BOD than in RNO.

The evening complex is a multi-protein complex composed of Early Flowering 3 (ELF3), Early Flowering 4 (ELF4) and Phytoclock 1 (PCL1 also known as LUX) that peaks at dusk. None of these proteins, had significant differences in transcript abundance between the two locations (Fig. 7; Additional file 12). The transcript abundance of ELF3 increased with sugar level and shortening of the day length (the higher sugar level comes later in the season and thus is at a shorter day length). ELF3, as part of the evening complex (EC), has direct physical interactions with PHYB, COP1 (Constitutive Photomorphogenic 1) and TOC1 [57] linking light and temperature signaling pathways directly with the circadian clock. It is interesting that most of the components of the clock showed significant differences in transcript abundance between BOD and RNO, except for the three proteins that make up the evening clock.

The transcript abundance profile of *PHYB* was similar in both BOD and RNO berries (Fig. 7), however the changes in transcript abundance with sugar level occurred in BOD berries at a lower sugar level. There was a gradual decline of *PHYB* transcript abundance with increasing sugar level until the last measurement at the fully mature stage, where there was a large increase in transcript abundance. A very similar profile is observed for Reveille 1 (*RVE1*). RVE1 promotes seed dormancy in Arabidopsis and PHYB interacts with RVE1 by inhibiting its expression [58]. PIF7 (Phytochrome Interacting Factor 7), interacts directly with PHYB to suppress PHYB protein levels [59]. Likewise, PIF7 activity is regulated by the circadian clock [60]. *PIF7* had higher transcript abundance in the BOD than that of RNO berries and generally increased with increasing sugar level. The transcript abundance of two of the other grape phytochromes (*PHYA* and *PHYE*) did not vary significantly between the two locations or at different sugar levels. *PHYC* had a higher transcript abundance in RNO berries and did not change much with different sugar levels. Many other light receptors (e.g. *CRY3* (cryptochrome 3), *FAR1* (far-red impaired response 1), *FRS5* (FAR1-related sequence 5), etc.) had higher transcript abundance in BOD berries (Additional file 13). Thus, light sensing through the circadian clock is a complicated process with multiple inputs.

RVE1 follows a circadian rhythm [61]. It behaves like a morning-phased transcription factor and binds to the EE element, but it is not clear if it is affected directly by the core clock (e.g. TOC1 or EC which repress other morning gene paralogs like *CCA1* and *LHY* (late elongated hypocotyl)) or through effects of PHYB or both. PHYB downregulates RVE1; RVE1 promotes auxin concentrations and decreases gibberellin (GA) concentrations [58]. Warmer night temperatures (as in BOD) cause more rapid reversion of the active form of PHYB to the inactive form [33] and thus may promote a higher expression/activity of RVE1. P_r_ (phytochrome in the red form, which is the physiologically inactive form) appears to accelerate the pace of the clock [55]. It is unclear what role phytochromes might have in seed and fruit development in grapes.

Very little is known about the effect of PHY on fruit development in general. In one tomato study, the fruit development of *phy* mutants was accelerated [62], suggesting that PHYB as a temperature/light sensor and a regulator of the circadian clock may influence fruit development. Carotenoid concentrations, but not sugar concentrations, also were affected in these mutants.

Photoperiod affects the transcript abundance of *PHYA* and *PHYB* in grape leaves [63]. In the present study, the transcript abundance of the majority of the photoreceptor genes in berry skins, including red, blue and UV light photoreceptors, had a higher transcript abundance in BOD berries (Additional file 13). It is unclear what the effect of PHYB and the circadian clock have on grape berry development. However, there were clear differences between the two locations; it seems likely that PHYB and the circadian clock are key grape berry sensors of the environment, affecting fruit development and composition.

### Transcript abundance of temperature-responsive genes

The grape berry transcriptome is sensitive to temperature [2, 3]. Temperature related genes were differentially expressed at the two locations in our study. The RNO berries were exposed to a much larger temperature differential between day and night than BOD berries and were also exposed to chilling temperatures in the early morning hours during the late stages of berry ripening (Table 1). The transcript abundance of some cold-responsive genes was higher in RNO berry skins than in BOD berry skins (Fig. 8), including *CBF1*.

*CBF1* transcript abundance is very sensitive to chilling temperatures; it is a master regulator of the cold regulon and improves plant cold tolerance [35, 64, 65]. PIF7 binds to the promoter of *CBF1*, inhibiting *CBF1* transcript abundance, linking phytochrome, the circadian clock and *CBF1* expression [60]. Our data are consistent with this model; transcript abundance of *PIF7* was higher and *CBF1* transcript abundance was lower in BOD berry skins than RNO berry skins (Fig. 7 and 7).

### Transcript abundance of dehydration and seed dormancy genes

ABA concentrations in plants increase in response to dehydration and ABA triggers a major signaling pathway involved in osmotic stress responses and seed development [66]. ABA concentrations only increase in the seed embryo near the end of seed development when the embryo dehydrates and goes into dormancy. ABA concentrations remain high to inhibit seed germination. The transcript abundance of ABA signaling genes such as *ABF2* and *SnRK2* (SNF1 related protein kinase 2) kinases increase after application of ABA to cell culture [67] and in response to dehydration [45] in leaves of Cabernet Sauvignon.

The data in this study are consistent with the hypothesis that BOD berries are riper at lower sugar levels. The ABA signaling genes in the berry skins had higher transcript abundance in BOD berries indicating that ABA concentrations were higher in BOD than RNO berries even though RNO berries were exposed to drier conditions (Table 1). ABA concentrations may be higher in the BOD berry skins based upon the higher transcript abundance of important ABA signaling and biosynthesis genes encoding ABF2, SnRK2 kinases and NCED6. We hypothesize that this would be seed derived ABA since water deficits were not apparent in BOD with the recent rainfall and high humidity. In contrast, *NCED3* and *NCED5* had higher transcript abundance in RNO berry skins, which might occur as the result of the very low humidity and large vapor pressure deficit (the vines were irrigated). The lower expression of *NCED6* in RNO berry skins may indicate that the seeds in the berry were more immature than the BOD berries. The higher expression of other seed development and dormancy genes (e.g. *RVE1*, *ARF2*, *ARF10*, etc.) in the berry skins support the argument that BOD berries (and seeds) matured at a lower sugar level than the RNO berries.

The ABA concentrations in the berry skins are a function of biosynthesis, catabolism, conjugation and transport. ABA in seeds increase as the seed matures and some of this ABA may be transported to the skin. In fact, a number of *ABCG40* genes, which encode ABA transporters, had higher transcript abundance in BOD berry skins than that in RNO (Additional file 2 and 14). Part of the ABA in skins may be transported from the seed and part of it might be derived from biosynthesis in the skins. *NCED6* transcript abundance in the skins was higher in BOD berries. Perhaps the transcript abundance of *NCED6* in the skin is regulated by the same signals as the embryo and reflects an increase in seed maturity. *AtNCED6* transcript abundance is not responsive to water deficit in Arabidopsis, but *AtNCED3* and *AtNCED5* are [43]. This is consistent with the higher *NCED3, NCED5* and *BAM1* transcript abundance in RNO berries (Additional file 14). Thus, there are complex responses of ABA metabolism and signaling. It would appear that there may be two different ABA pathways affecting ABA concentrations and signaling: one involved with embryo development and one involved with the water status in the skins.

Auxin is also involved with ABA signaling during the late stages of embryo development in the seeds. Auxin signaling responses are complex. ABF5 is an auxin receptor that degrades Aux/IAA proteins, which are repressors of ARF transcriptional activity [68]. Thus, a rise in auxin concentration releases Aux/IAA repression of ARF transcription factors, activating auxin signaling. In the berry skins, there was a diversity of transcriptional responses of *Aux/IAA* and *ARF* genes in the two locations, some with increased transcript abundance and others with decreased transcript abundance. As with ABA signaling, there may be multiple auxin signaling pathways operating simultaneously.

One pathway appears to involve seed dormancy. *ARF2* had a higher transcript abundance in BOD berries. ARF2 promotes dormancy through the ABA signaling pathway [69]. This is consistent with the hypothesis that BOD berries reach maturity at a lower sugar level than RNO berries.

### Transcript abundance of biotic stress genes

Grapevines have very dynamic gene expression responses to pathogens [70, 71]. The top 150 DEGs for BOD berries were highly enriched with biotic stress genes. The higher rainfall and high relative humidity in BOD would make moist conditions suitable for pathogenic fungi to grow. We detected a much higher transcript abundance of powdery mildew-responsive genes in BOD berries and this may be connected to a higher transcript abundance of ethylene and phenylpropanoid genes as part of a defense response. The transcript abundance profiles of some of these genes (e.g. *PR10*, *PAL1*, *STS10*, *ACS6*, and *ERF2*; see Figs. 5, 8, 9 and Additional file 11) are remarkably similar.

Increased ethylene signaling in grapevines has been associated with powdery mildew infection and phenylpropanoid metabolism and appears to provide plant protection against the fungus [72, 73]. Genes involved with phenylpropanoid metabolism, especially *PAL* and *STS* genes, appear to be quite sensitive to multiple stresses in the environment [74]. In Arabidopsis there are four *PAL* genes [75]. These *PAL* genes appear to be involved with flavonoid biosynthesis and pathogen resistance in Arabidopsis. Ten different *PAL1* and two *PAL2* orthologs had higher transcript abundance in BOD berry skins; many *STS* genes also had a higher transcript abundance in BOD berry skins (Additional file 11). Stilbenes are phytoalexins and provide pathogen resistance in grapes and *STS* genes are strongly induced by pathogens [70]. Thus, the higher transcript abundance of powdery mildew genes may be associated with the higher transcript abundance of genes in the ethylene and phenylpropanoid pathways.

### Transcript abundance of iron homeostasis genes

The transcript abundance of a number of iron homeostasis genes were significantly different in the two locations (Fig. 11) and there was a difference in soil available iron concentrations in the two locations. However, iron uptake and transport in plants is complicated depending on multiple factors, such as pH, soil redox state, organic matter composition, solubility in the phloem, etc. Thus, it is impossible to predict iron concentrations in the berry without direct measurements. The roles of these genes in iron homeostasis and plant physiological functions are diverse. Iron supply can affect anthocyanin concentrations and the transcript abundance of genes in the phenylpropanoid pathway in Cabernet Sauvignon berry skins [76]. One of the DEGs, *SIA1*, is located in the chloroplast in Arabidopsis and appears to function in plastoglobule formation and iron homeostasis signaling in concert with *ATH13* (also known as OSA1) [77]. Another DEG, *YSL3*, is involved in iron transport [78]. It acts in the SA signaling pathway and appears to be involved in defense responses to pathogens. It also functions in iron transport into seeds [79]. FER1 is one of a family of ferritins (iron-binding proteins) found in Arabidopsis [80]. VIT1 and NRAMP3 are vacuolar iron transporters [81] and are also involved in iron storage in seeds. Other DEGs are also responsive to iron supply. *IREG3* (also known as *MAR1*) appears to be involved in iron transport in plastids; its transcript abundance increases with increasing iron concentrations [82]. *ABCI8* is an iron-stimulated ATPase located in the chloroplast that functions in iron homeostasis [83].

It is unclear what specific roles these iron homeostasis genes are playing in grape berry skins, but they appear to be involved in iron storage in seeds and protection against oxidative stress responses [80, 81]. One possible explanation for the transcript abundance profiles in the BOD and RNO berry skins is that ferritins are known to bind iron and are thought to reduce the free iron concentrations in the chloroplast, thus, reducing ROS production that is caused by the Fenton reaction [80]. As chloroplasts senesce during berry ripening, iron concentrations may rise as a result of the catabolism of iron-containing proteins in the thylakoid membranes; thus, berry skins may need higher concentrations of ferritins to keep free iron concentrations low. This might explain the increase in ferritin transcript abundance with increasing sugar levels.

Most soils contain 2 to 5% iron including available and unavailable iron; soils with 15 and 25 µg g^-1^ of available iron are considered moderate for grapevines [84], but soils with higher concentrations are not considered toxic. Therefore, for both soils in this study, iron concentrations can be considered to be very high but not toxic. The higher available iron concentrations in the BOD vineyard may be associated with the wetter conditions (more reductive conditions) and the lower soil pH.

### Environmental influences on transcript abundance in other studies

Other researchers using Omics approaches have identified environmental factors that influence grape berry transcript abundance and metabolites. One study investigated the differences in transcript abundance in berries of Corvina (a black-skinned grape cultivar that makes red wine) in 11 different vineyards within the same region over three years [85]. They determined that approximately 18% of the berry transcript abundance was affected by the environment. Climate had an overwhelming effect but viticultural practices were also significant. Phenylpropanoid metabolism was very sensitive to the environment and *PAL* transcript abundance was associated with *STS* transcript abundance.

In another study of a white grape cultivar, Garganega, berries were analyzed by transcriptomic and metabolomic approaches [86]. Berries were selected from vineyards at different altitudes and soil types. Again, phenylpropanoid metabolism was strongly influenced by the environment. Carotenoid and terpenoid metabolism were influenced as well.

Two studies investigated the grape berry transcriptomes during the ripening phase in two different regions of China, a dry region in Western China and a wet region in Eastern China [87, 88]. These two locations mirror some of the differences in our conditions in our study, namely moisture, light and elevation, although the dry China western region has higher night temperatures and more rainfall than the very dry RNO location. In the Cabernet Sauvignon study [87], they compared the berry transcriptomes (with seeds removed) from the two regions at three different stages: pea size, veraison and maturity. The TSS at maturity was slightly below 20°Brix. Similar to our study, the response to stimulus, phenylpropanoid and diterpenoid metabolism GO categories were highly enriched in mature berries between the two locations. Differences in the transcript abundance of NCED and PR proteins were also noted. Like in our study, the authors associated the transcript abundance of these proteins to the dry (drought response) and wet (pathogen defense) locations, respectively.

In the second study comparing these two regions in China [88], the effects of the environment on the metabolome and transcriptome of Muscat Blanc à Petits Grains berries were investigated over two seasons; specifically, terpenoid metabolism was targeted. Like in our study, the transcripts in terpenoid were in higher abundance in the wetter location. The transcript abundances were correlated with terpenoid concentrations and a coexpression network was constructed. A specific set of candidate regulatory genes were identified including some terpene synthases (TPS14), glycosyl transferases and 1-hydroxy-2-methyl-2-butenyl 4-diphosphate reductase (HDR). We examined the transcript abundance of some of these candidate genes in our own data but did not find significant differences between our two locations. The contrasting results between our study and Wen et al. (2015) could be for a variety of reasons such as different cultivar responses, berry versus skin samples, or different environmental conditions that affect terpenoid production.

Terpenoid metabolism is influenced by the microclimate [89] and is involved in plant defense responses to pathogens and insects [29, 90]. Light exposure to Sauvignon Blanc grapes (a white grape cultivar) was manipulated by removing adjacent leaves without any detectable differences in berry temperatures [89]. Increased light exposure increased specific carotenoid and terpene concentrations in the berry. The responses of carotenoid and terpenoid production to temperature are less clear. Some effect of temperature was associated with carotenoid and terpenoid production, but to a lesser extent than light [89]. Higher concentrations of rotundone, a sesquiterpene, have been associated with cooler temperatures [91]. Water deficit can also alter carotenoid and terpenoid metabolism in grapes [11, 92]. Terpenes can act as signals for insect attacks and attract insect predators [90]. Thus, terpenoid metabolism is highly sensitive to the environment and influenced by many factors.

In contrast to these studies, excess light and heat can affect transcript abundance and damage berry quality. In addition to a higher rate of malate catabolism, anthocyanin concentrations and some of the transcript abundances associated with them are decreased as well [93, 94].

### Temperature effects on berry maturity and total soluble solids

BOD berries reached maturity at a lower °Brix level than RNO berries; the cause is likely to be the warmer days and cooler nights in RNO. Higher day temperature may increase photosynthesis and sugar transport and cooler night temperatures may reduce fruit respiration. °Brix or TSS approximates the % sugar in a berry and is a reliable marker of berry maturity in any given location [95]; however, TSS is an unreliable marker of berry maturity when comparing grapes from very different climates. The differences in TSS between BOD and RNO are consistent with other studies on the temperature effects on berry development. Indirect studies have associated gradual warming over the last century to accelerated phenology and increased sugar concentrations in the grape berries [96–99]. Increasing temperature can accelerate metabolism, including sugar biosynthesis and transport, but the increase in metabolism is not uniform. For example, the increase in anthocyanin concentration during the ripening phase is not affected as much as the increase in sugar concentration [100]. These responses vary with the cultivar [97], complicating this kind of analysis even further.

Direct studies of temperature effects on Cabernet Sauvignon berry composition also are consistent with our data. In one study, the composition of Cabernet Sauvignon berries was altered substantially for vines grown in phytotrons at 20 or 30°C temperatures (temperatures that are very similar to the BOD and RNO temperatures occurring in the present study) [101]. Cooler temperatures promoted anthocyanin development and malate concentrations (inhibited malate catabolism) and higher temperatures promoted TSS (°Brix) and proline concentrations [101]. In a second study, vines were grown at 20 or 30°C day temperatures with night temperatures 5°C cooler than the day [102]. In this study, higher temperatures increased berry volume and veraison started earlier by about 3 to 4 weeks [102]. The authors concluded that warmer temperatures hastened berry development. In a third study, Cabernet Sauvignon berry composition was affected in a similar manner by soil temperatures that differed by 13°C [103].

TSS concentrations are also affected by light and the vine water status. Light is generally not a factor because there is usually a large enough leaf area and sufficient light levels to saturate this source to sink relationship [104, 105]. Sun-exposed Cabernet Sauvignon berries in the vineyard had higher TSS than shaded berries [104]. This sunlight effect was attributed largely to an increase in berry temperature rather than an increase in the fluence rate per se. A higher grapevine water status results in larger berry size and lower sugar concentrations [106] and water deficit is known to increase sugar concentrations in Cabernet Sauvignon [11]. However, temperature is thought to have the largest effect on sugar concentrations [16].

Other transcriptomic data in the present study indicated that BOD berries were more mature at a lower sugar level than RNO berries. These included the transcript abundance profiles of genes involved in autophagy, auxin and ABA signaling, iron homeostasis and seed development. Many of these DEGs had an accelerated rate of change in BOD berries. While these transcripts are in the skins, they may be influenced by signals coming from the seed. In addition, there was a higher transcript abundance for most genes involved with the circadian clock in BOD berries. PHYB can regulate the circadian clock [55] and PHYB activity is very sensitive to night temperatures (BOD had higher night temperatures); PHYB reversion is accelerated to the inactive form at warmer temperatures [33]. The inactivity of phytochrome promotes the expression of RVE1, which promotes auxin concentrations and seed dormancy [58]. Thus, all things considered, it is likely that temperature and/or the temperature differentials between day and night significantly contributed to the differences in the rate of berry development and sugar accumulation in the two locations.

### Are there reliable markers to harvest berries at maturity?

Determining maturity of grapes is a difficult and error prone process. Reliable markers could aid in the decision of when to harvest the grapes. “Optimum” maturity is a judgement call and will ultimately depend on the winemaker’s or grower’s specific goals or preferences. A combination of empirical factors can be utilized including °Brix, total acidity, berry tasting in the mouth for aroma and tannins, seed color, etc. °Brix or total soluble solids by itself may not be the best marker for berry ripening as it appears to be uncoupled from berry maturity by temperature. Phenylpropanoid metabolism, including anthocyanin metabolism, is also highly sensitive to both abiotic and biotic stresses and may not be a good indicator of full maturity. Thus, color may not be a good indicator either. Specific developmental signals from the seed or embryo, such as those involved with auxin and ABA signaling, may provide more reliable markers for berry ripening in diverse environments, but will not be useful in seedless grapes. Aromatic compounds may also be reliable markers but they will need to be generic, developmental markers that are not influenced by the environment. This study revealed many genes that are <u>not</u> reliable markers because they were expressed differently in different environments. One candidate marker that is noteworthy is ATG18G (g071260). Its transcript abundance increased and was relatively linear with increasing °Brix and these trends were offset at the two locations relative to their level of putative fruit maturity (Fig. 4). ATG18G is required for the autophagy process [107] and maybe important during the fruit ripening phase. It was found to be a hub gene in a gene subnetwork associated with fruit ripening and chloroplast degradation [4]. Further testing will be required to know if it is essential for fruit ripening and whether its transcript abundance is influenced by abiotic and biotic stresses in grape berry skins.

### Conclusions

The ultimate function of a fruit is to produce fully mature seeds in order to reproduce another generation of plants. Berry ripening involves complex signaling that tells other organisms when the fruit is ready for consumption and seed dispersal. In this study, we tested and confirmed the hypothesis that the transcript abundance in grape skins differed in two different locations with different environments. Another goal was to distinguish transcripts with common profiles from differentially expressed genes. The observations made in this study provide lists of such genes and generated a large number of hypotheses to be tested in the future. WGCNA was particularly powerful and enhanced our analyses. It is clear that berry ripening may respond to a multitude of factors both before and during the late stages of berry ripening. The transcriptomic analysis in the late stages of berry ripening in this study indicated that transcript abundance was very dynamic and possibly influenced by a number of environmental and internal (e.g. seed signals) factors. We could discern these influences using functional analysis of the genes and GO enrichment analysis of the transcriptomic datasets. Temperature, light, moisture and the local microbiome were major factors that may have contributed to the transcript abundance profiles of the berry skins. While earlier fruit development stages clearly have an effect on fruit composition, the dynamic changes in transcript abundance in the late stages of berry ripening indicated that berries still have a “sense of place”; it appears that the environment continues to affect berry metabolite composition, which may impact berry flavor, color and other quality-related traits. Future studies are required to follow up on these observations. It appears that fruit ripening is very malleable. Manipulation of the canopy (time and intensity of leaf removal at different locations on the plant) may offer a powerful lever to adjust gene expression and berry composition, since these parameters are strongly affected by light and temperature.

## Methods

### Plant materials

Grapevines in Reno (*Vitis vinifera* L. cv. Cabernet Sauvignon clone 8) were grown on their own roots at the Valley Road Experimental Vineyard at the Nevada Agricultural Experiment Station on the campus of the University of Nevada, Reno. Research approval was obtained by Grant R. Cramer from the Nevada Agricultural Experiment Station and the University of Nevada, Reno. The grapevines were originally obtained as certified material from Inland Desert Nursery, Benton, City, Washington, USA. The grapes were harvested between September 10 and October 2012 depending on maturity. Maturity was assessed using a digital refractometer (HI 96811, Hanna Instruments, Woonsocket, RI, USA) to measure soluble solids (°Brix) that are mostly made up of sugars. Berry clusters were collected between 11.00 h and 13.00 h (near solar noon) in an attempt to minimize temporal transcriptional response variations caused by the circadian clock. At harvest, individual berry °Brix levels were determined with a digital refractometer. Separated berry skins were placed into 50 mL centrifuge tubes in liquid nitrogen according to sugar level (1 ± 0.5 °Brix increments; 19 to 27 °Brix). In this way berries were collected over many days from multiple clusters from multiple vines from 3 different individually drip-irrigated blocks in the vineyard. Stem-water potentials were measured weekly to determine irrigation levels that would maintain the water status of the vines. The vines were regularly sprayed for fungal prevention. Each block in the vineyard was considered an experimental replicate. Soil testing was performed by A & L Western Agricultural Laboratories (Modesto, CA, USA).

Cabernet Sauvignon clone CA33 412 grapevines in Bordeaux were grown on SO4 rootstock at the VitAdapt parcel on the Institut National de Recherche Agronomique (INRA) research station in Villenave d’Ornon, in the Aquitaine region of France. There were 5 replicate blocks within this site to mitigate soil variation. Additional details of the VitAdapt project can be found in [108]. Berries for the three replicates were collected at the 14:00 h of the day. Environmental conditions and variables of the Reno and BOD vineyards are listed in Table 1. The vines were regularly sprayed for fungal prevention. Soil testing was performed by Aurea Agroscience (Blanquefort, France).

### RNA extraction

Three experimental replicates from each cultivar at 20, 22, 24 and 26 °Brix were used for sequencing. In RNO, total RNA was extracted from approximately 250 mg of frozen, finely ground, skin tissue using a modified CTAB extraction protocol followed by an additional on-column DNase digestion using a Qiagen RNeasy Mini Kit (Qiagen, Valencia, CA, USA) as in [109]. RNA quality and quantity were assessed with a Nanodrop ND-1000 spectrophotometer (ThermoFisher Scientific, Waltham, MA, USA) and an Agilent 2100 Bioanalyzer (Agilent Technologies, Santa Clara, CA, USA). In BOD, total RNA was extracted according to Reid et al. [110] from 1 g of frozen, finely ground berry skins. Traces of genomic DNA were removed by a DNAse I treatment according to the manufacturer’s instructions (Ambion TURBO DNA-free DNase, Life Technologies). The RNA was quantified using a Nanodrop 2000c spectrophotometer (Thermo Scientific) and their integrity was checked on an 1.8% agarose gel.

### RNA-Seq library preparation and sequencing

Cabernet Sauvignon 50 bp single-end, barcoded libraries were constructed and sequenced by the Neuroscience Genomics Core at the University of California, Los Angeles for the RNO samples and by the Genome Center at the University of California, Davis for BOD samples using Illumina TruSeq RNA library prep kits (Illumina Inc., San Diego, CA, USA) according to the manufacturer’s instructions. The barcoded libraries were pooled, multiplexed, and were sequenced using Illumina TruSeq chemistry (version 3.0) on a HiSeq2000 sequencer (Illumina Inc., San Diego, CA, USA).

### Transcript abundance and statistical analysis

The single-end sequence fragments (reads) generated by Illumina sequencing were base-called and demultiplexed. The fastq files generated were inspected for sequence quality and contamination using FastQC [111]. Illumina adapters were removed from sequences with Trimommatic [112] version 0.36. The transcript abundance was estimated from the trimmed fastq files using Salmon [24] version 0.14.1 using --gcBias, --seqBias, --fldMean=50, --fldSD=1, --validateMappings --rangeFactorizationBins 4. An augmented hybrid fasta file was built from the *Vitis vinifera* cv. Cabernet Sauvignon genome [26] using generateDecoyTranscriptome.sh from salmontools. This file was used to build the index file used for the quantification with a k-mer size of 15. The salmon output (quant files) was imported into DESeq2 [27] version 1.22.2 using tximport version 1.10.1 [113] for determination of differentially expressed genes (DEGs).

### Coexpression network analysis

A coexpression gene network analysis was performed using WGCNA version 1.68 [114, 115] using all the libraries for each location. Prior to this analysis, low-expressed genes were removed with a minimum threshold of 10 counts in all the libraries. Counts data were transformed using the function varianceStabilizingTransformation of the package DESeq2. The resulting set of counts was used for network construction and module detection using the function blockwiseModules. An adjacency matrix was created by calculating the biweight mid-correlation raised to a power β of 8 and the maxPoutliers parameter set to 0.05. The subsequent Topological Overlap Matrix (TOM) was used for module detection using the DynamicTreecut algorithm with a minimal module size of 30 and a branch merge cut height of 0.25. The module eigengenes were used to evaluate the association between the resulting modules (22) and the experimental traits (Brix degrees and locations).

### Gene functional annotation additions

Gene models of the Cabernet Sauvignon annotation were searched against different protein databases with the blastx function of the DIAMOND version 0.9.19 software [116] using default parameters and reporting alignments in the 1% range of the top alignment score. For each gene model, the best blast hit was kept (1-to-1) and reported in addition of the current annotation. For multiple hits with the same score (1-to-many), the first hit was kept as the representative result but the other hits are still accessible. The databases used were Araport11, release 06.17.16, [117] and the *Vitis vinifera* IGGP 12X from EnsemblPlants 38, a part of EnsemblGenomes [118]. The corresponding gene annotations were obtained from the Araport11 gff file (release 06.22.16), TAIR10 functional descriptions (release 01.16.13) and a manually curated and actualized grapevine V1 annotation of PN40024.

### Functional enrichment of GO categories

Gene Ontology (GO) enrichment was performed using topGO version 2.34.0 [119]. Enriched functional categories with an FDR adjusted p-value > 0.01 after the Fisher’s test were filtered for further analysis. For gene ontology (GO) categories assignments, the GO already present in the Cabernet Sauvignon annotation file [25] were combined with the previously manually curated GO annotations of the PN40024 V1 gene models. The GO from P40024 were attributed to the Cabernet Sauvignon gene model if the blast hit was presenting a percentage of identity greater than 95 % as well as an alignment representing more than 95% of the length of both the query and the subject.

## Abbreviations

ABA: ABscisic Acid
ABCG: Adenosine triphosphate Binding Casette G
ABF: Abscisic acid responsive elements Binding Factor
ABS: Abnormal Shoot
ACO1: ACc Oxidase 1
ACOL: ACc Oxidase-Like
APG9: AutoPhaGy 9
ARAC1: Arabidopsis RAC-like 1
ARF: Auxin Response Factor
ATG: AuTophaGy
ATH: Arabidopsis Thaliana abc2 Homolog
BAM: Barley Any Meristem
BCAT: Branched-Chain-amino-acid AminoTransferase
BOD: BOrDeaux
bp: base pair BSMT1: Benzoate/Salicylate MethylTransferase 1
CBF1: C-repeat/DRE Binding Factor 1
CCA1: Circadian Clock Associated 1
CLE: CLavata3/Esr-related
COP1: COnstitutive Photomorphogenic 1
CML: CalModulin-Like
CRY3: CRYptochrome 3
cv: cultivar
CYSB: cystatin B
DEG(s): Differentially Expressed Gene(s)
DHS: 3-deoxy-D-arabino-Heptulosonate 7-phosphate Synthase
DMAPP: DiMethylAllyl diPhosPhate
DUF642: Domain of Unknown Function 642
DXR: 1-deoxy-D-Xylulose 5-phosphate Reductoisomerase
EC: Evening Complex
EE: Evening Element
EIN: Ethylene INsensitive
ELF: EarLy Flowering
ERF: Ethylene Response Factor
EXL2: EXordium Like 2
FAR1: Far-Red impaired Response 1
FDR: False Discovery Rate
FER: FERrittin
FOP1: FOlded Petal 1
FRS: FAR1-Related Sequence
GH3: GH3 family protein
GIDB1: Gibberellic acid Insensitive Dwarf1B
GO: Gene Ontology
HAD: HaloAcid Dehalogenase-like hydrolase protein
HB12: HomeoBox 12
HDS: 4-Hydroxy-3-methylbut-2-en-1-yl Diphosphate Synthase
IAA: Indole Acetic Acid
IDD14: InDeterminate-Domain 14
IPP: IsoPentenyl Pyrophosphate
IREG: Iron REGulated
JAR: JAsmonate Resistant
MAT3: Methionine AdenosylTransferase 3
LHY: Late elongated Hypocotyl
MEP: MethylErythritol 4-Phosphate
MLA: intracellular MiLdew A
NAC073: NAC domain containing protein 73
NCBI: National Center for Biotechnology Information
NCED: Nine-Cis Epoxycarotenoid Dioxygenase
NRAMP3: Natural Resistance-Associated Macrophage Protein 3
PAL: PhenylAlanine Lyase
PCL1: PhytoCLock 1
PHY: PHYtochrome
PIF: Phytochrome Interacting Factor
PR: Pathogenesis Related
PRR: Psuedo-Response Regulator
PSY: Phytoene SYnthase
RNO: ReNO
ROS: Reactive Oxygen Species
RVE1: ReVeillE 1
SA: Salicylic Acid
SIA: Salt Induced Abc kinase
SnRK: SNF1 Related protein Kinase
STS: STilbene Synthase
SULTR: SULfate TRansporter
TAIR: The Arabidopsis Information Resource
TAT: TyrosineAmino Transferase
TOC: Timing Of Cab expression
TPPD: Trehalose Phosphate Phosphatase D
TPL: ToPLess
TPM: Transcripts Per Million
TPS: TerPene Synthase
TSS: Total Soluble Solids
VIT: Vacuolar Iron Transporter
WGCNA: Weighted Gene Co-expression Network Analysis
XTH: Xyloglucan endoTransglucosylase/Hydrolase
YSL: Yellow Stripe-Like

## Declarations

### Ethics

Not applicable.

### Consent to publish

Not applicable.

### Competing interests

The authors declare that they have no competing interests. The first author is a section editor of BMC Plant Biology.

### Authors’ contributions

GRC, SG, and AD-I designed the experiments. RG and AH collected berries, analyzed the °Brix levels and extracted the RNA. The original data were analyzed with the PN40024 genome annotation by GRC, but NC updated the analysis with the new Cabernet Sauvignon genome annotation, performed the WGCNA and assisted with estimation of the TPMs. GRC processed and analyzed the data and wrote the body of the paper. All authors reviewed, edited and approved the final version of the manuscript.

### Availability of supporting data

RNA-Seq data were deposited with the Sequence Read Archive (SRA) database at the National Center for Biotechnology Information (NCBI) with the SRA accession number PRJNA260535 [120] for the RNO data and SRP149949 for the BOD data.

## Acknowledgements

We would like to thank Aude Habran for her assistance in collecting berry skins and RNA extraction of BOD berries and thank the many UNR Biochemistry senior thesis undergraduate students that assisted in berry separation in the RNO berries.

## Funding

This research was supported by the University of Nevada, Reno, Agriculture Experimental Station with the Multi-state USDA-Hatch Grant NEV00383A to GRC.

## Supplemental Information

**Additional file 1.** Log_2_ transcripts per million (TPM) to genes uniquely mapped to the V1 Cabernet Sauvignon genome.

**Additional file 2.** Differentially expresses genes (DEGs) determined by DESeq2 for berry skin samples at 22°Brix from BOD and RNO. **Additional file 3.** Statistical results from gene set enrichment analysis of the DEGs using topGO.

**Additional file 4.** Image of the top 25 connected GO categories in the topGO network of the DEGs in Additional file 3.

**Additional file 5.** Common genes between BOD and RNO from transcriptomic approaches 2 and 3.

**Additional file 6.** Module membership (MM) of all filtered transcripts as defined by WGCNA. Values are the kME (module eigengene connectivity).

**Additional file 7.** Heatmap correlation of berry traits (°Brix level, BOD, RNO) of each of 19 gene modules. Gene modules were identified by a color name (MMcolornumber) as assigned by the WGCNA R package. Values in each heatmap block are the correlation (left value) and p-value (in parentheses) of the module with the berry trait.

**Additional file 8.** topGO analysis of all modules with genes having a kME >0.80. The results for each module are on a separate tab within the file.

**Additional file 9.** topGO analysis of the top 400 genes with the greatest differential expression between high and low °Brix samples in BOD berry skins.

**Additional file 10.** topGO analysis of the top 400 genes with the greatest differential expression between high and low °Brix samples in RNO berry skins.

**Additional file 11.** Representative examples of transcript profiles of some stilbene synthase (STS) genes that were differentially expressed.

**Additional file 12**. Transcript abundance of circadian clock genes from BOD and RNO berry skins. The data are placed on a circadian clock model derived from [4]. Lines in the model represent known interactions between genes; red arrows are positive interactions, black lines are negative interactions, and blue lines indicate direct physical interactions but the direction, positive or negative, is unknown. No lines indicate that there are no known interactions at this time. Transcript profiles outlined in red highlight significantly higher transcript abundance for the BOD berries.

**Additional file 13**. A model of the peripheral genes including light sensing genes that interact with core circadian clock genes in BOD and RNO berry skins. Lines represent gene interactions as described in Additional file 12. Red and blue lightning bolts represent the reception of their respective light wavelengths for each gene symbol. Transcript profiles outlined in red highlight significantly higher transcript abundance for the BOD berries.

**Additional file 14**. Transcript profiles of ABA biosynthesis and signaling genes that are differentially expressed between BOD and RNO berry skins.

## Notes

#### Summary of Updates

This version of the manuscript has been updated to incorporate the latest annotation of the Cabernet Sauvignon in place of the previously analyzed PN40024 genome and to add additional analyses to the manuscript at the request of reviewers.

