## Supplementary figures and images for "A Sense of Place: Transcriptomics Identifies Environmental Signatures in Cabernet Sauvignon Berry Skins in the Late Stages of Ripening"

### Additional File 4

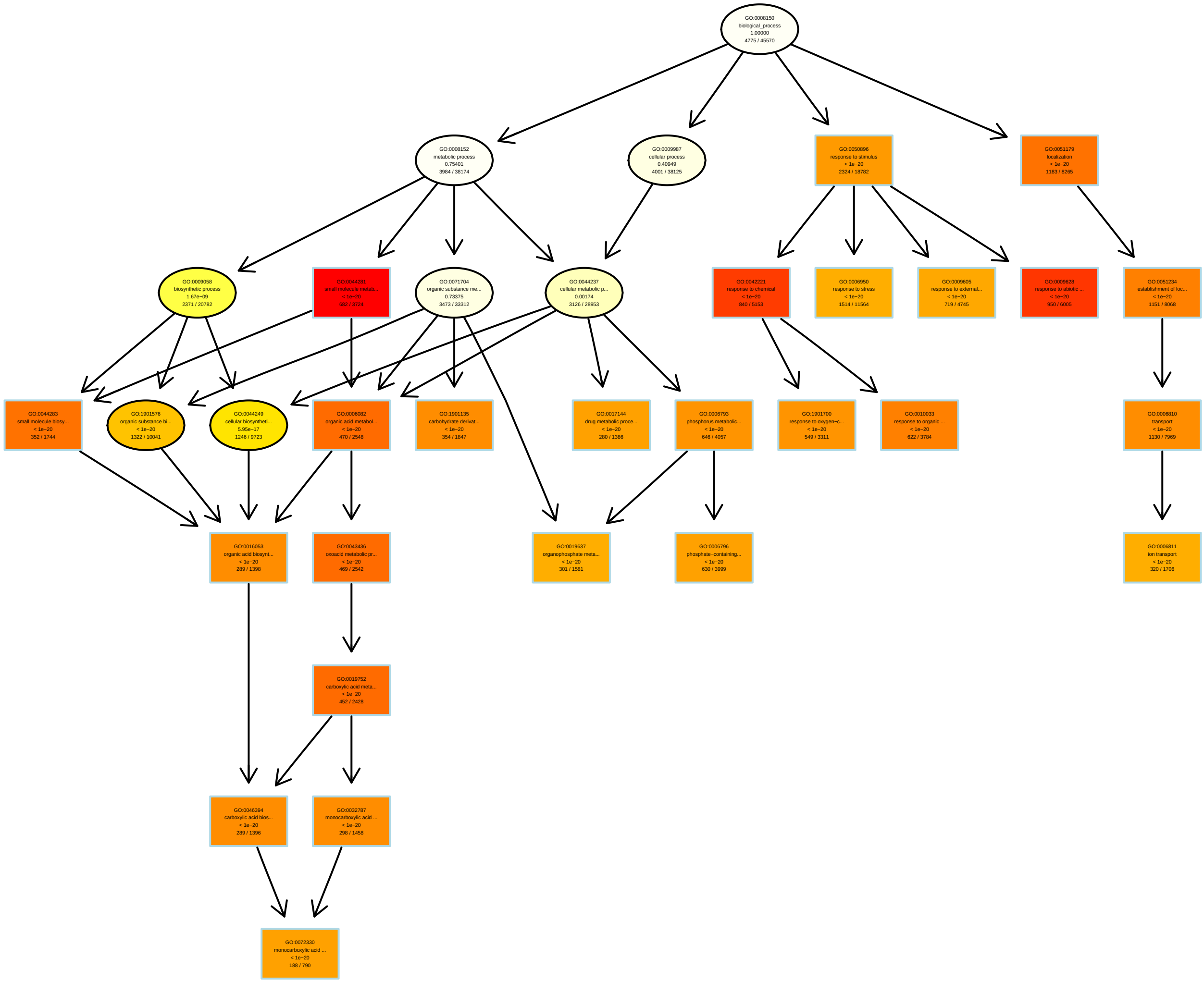

### Additional File 7

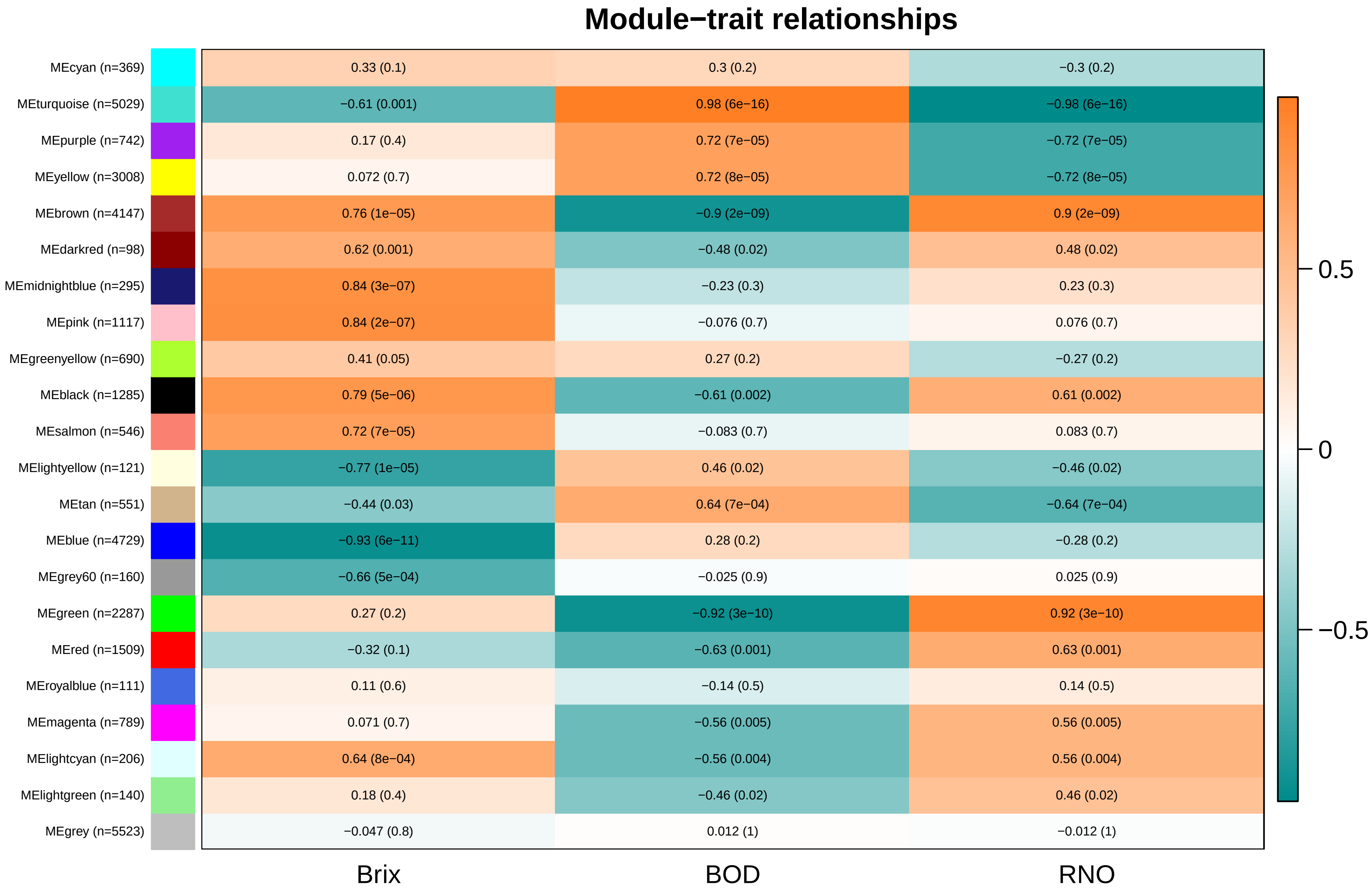

### Additional File 11

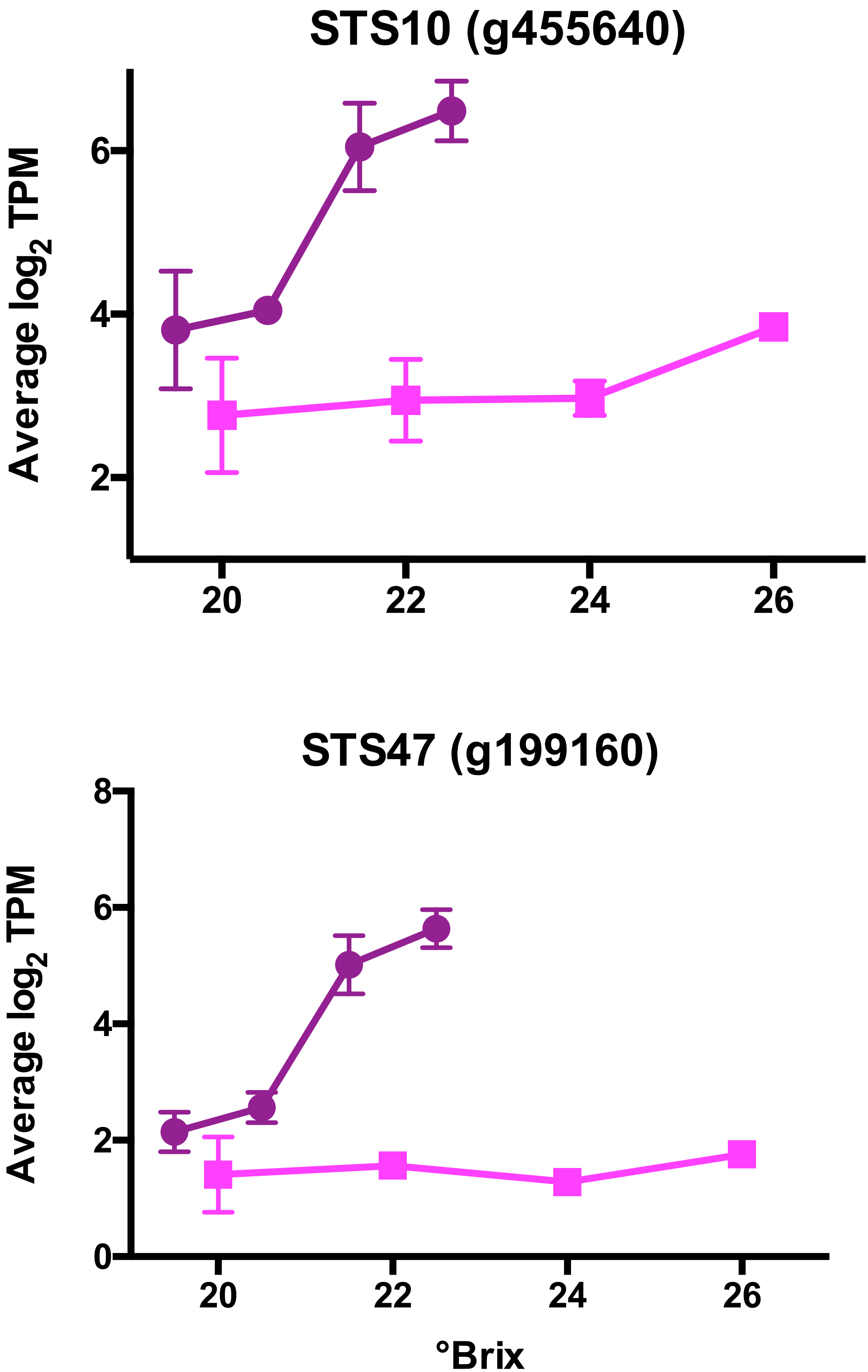

### Additional File 12

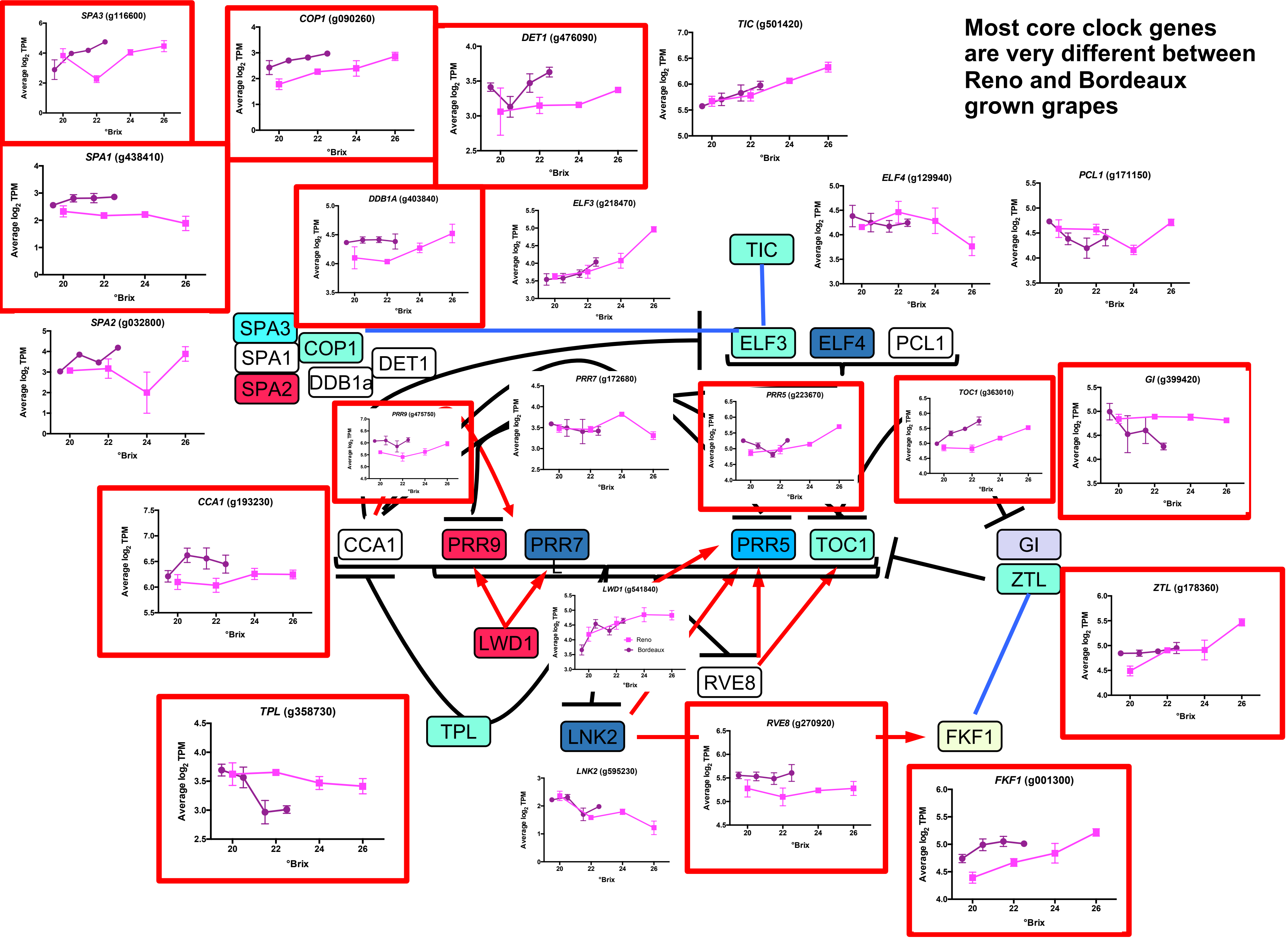

### Additional File 13

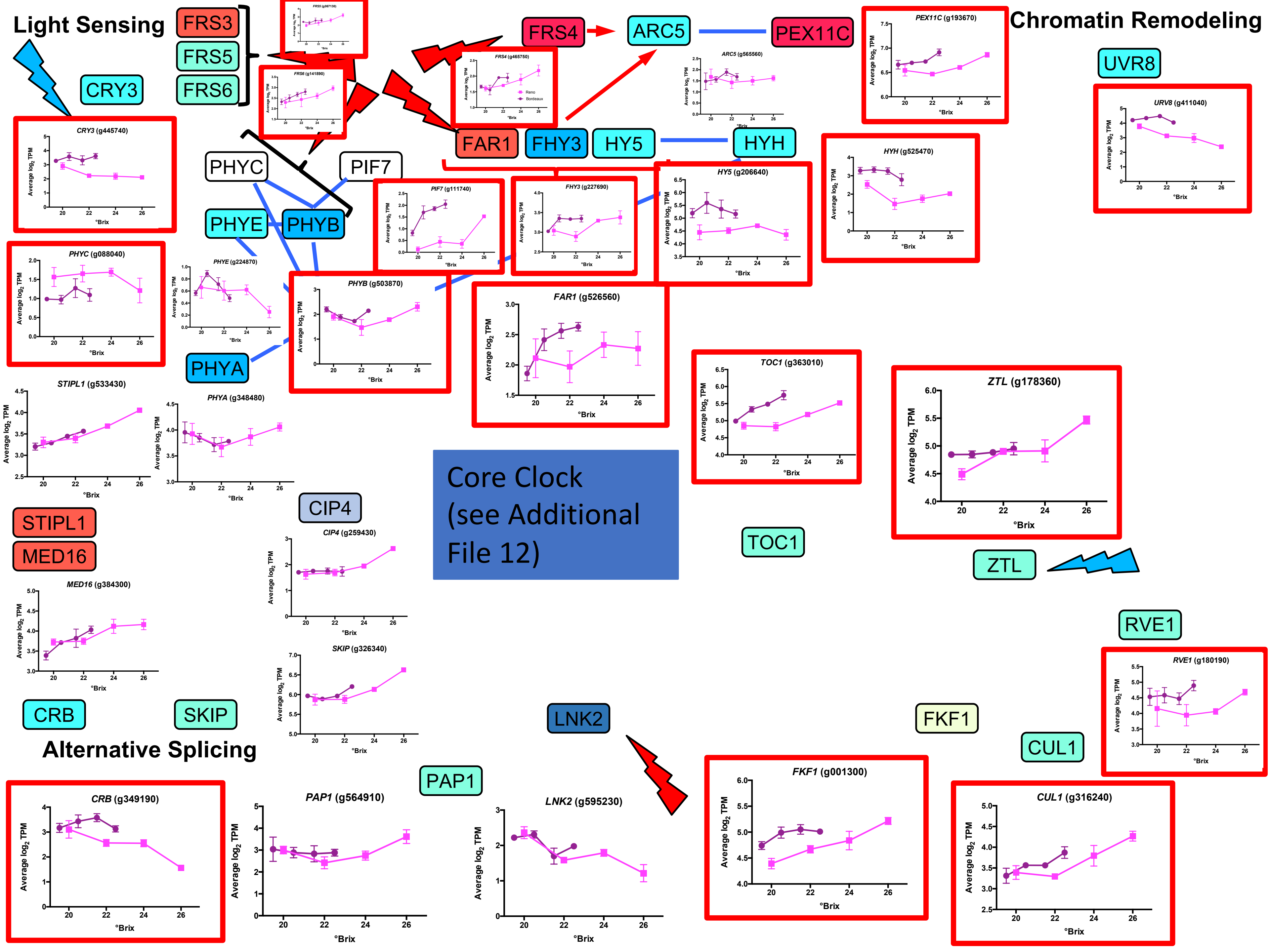

### Additional File 14

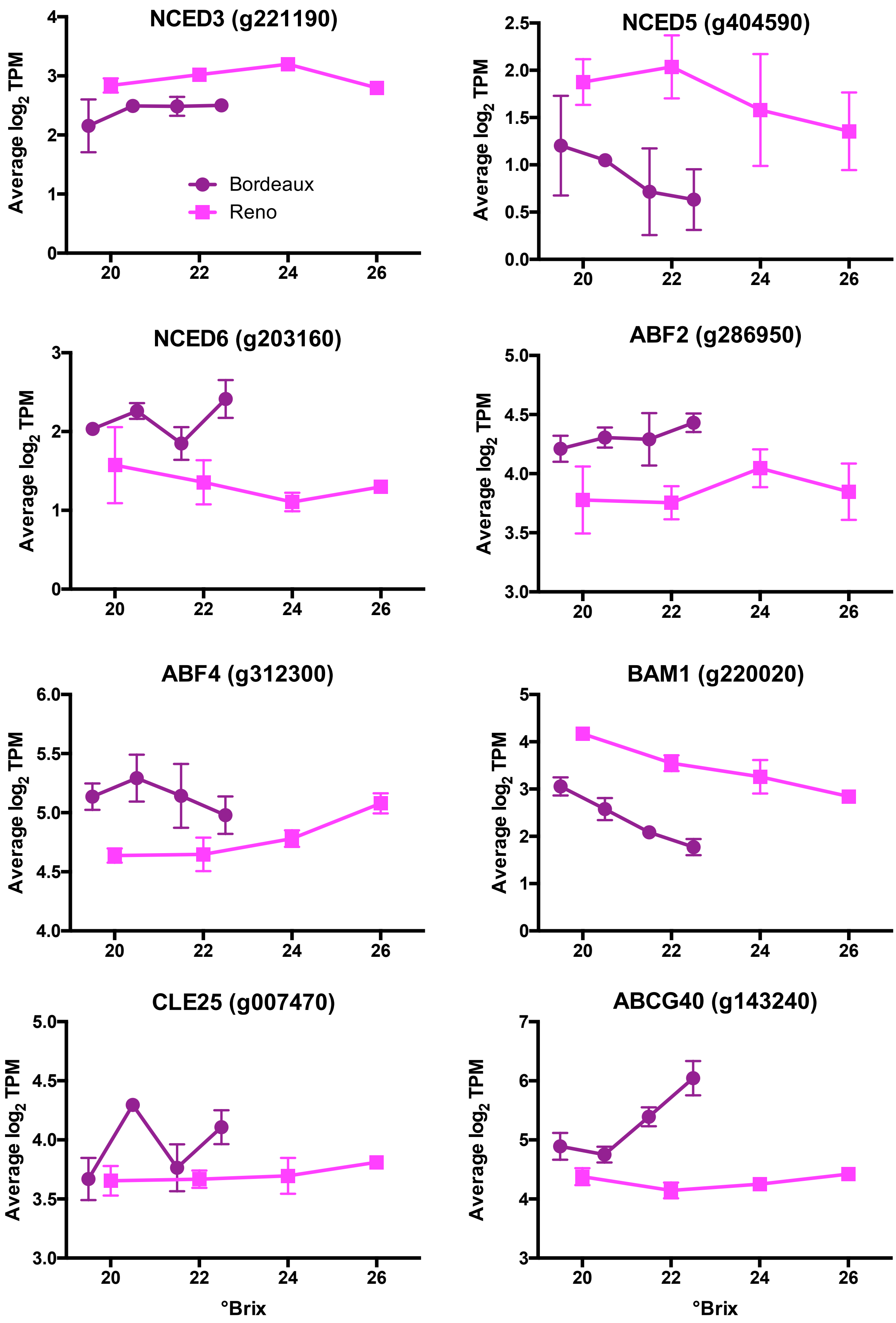
